# Responses of Model Cortical Neurons to Temporal Interference Stimulation and Related Transcranial Alternating Current Stimulation Modalities

**DOI:** 10.1101/2022.05.04.490540

**Authors:** Boshuo Wang, Aman S. Aberra, Warren M. Grill, Angel V. Peterchev

**Author notes:** A. S. Aberra is currently with the Department of Biological Sciences, Dartmouth College, Hanover, NH 03755, USA.

## Abstract

**Objective:** Temporal interference stimulation (TIS) was proposed as a non-invasive, focal, and steerable deep brain stimulation method. However, the mechanisms underlying experimentally-observed suprathreshold TIS effects are unknown, and prior simulation studies had limitations in the representations of the TIS electric field (E-field) and cerebral neurons. We examined the E-field and neural response characteristics for TIS and related transcranial alternating current stimulation modalities.

**Approach:** Using the uniform-field approximation, we simulated a range of stimulation parameters in biophysically realistic model cortical neurons, including different orientations, frequencies, amplitude ratios, amplitude modulation, and phase difference of the E-fields, and obtained thresholds for both activation and conduction block.

**Main results:** For two E-fields with similar amplitudes (representative of E-field distributions at the target region), TIS generated an amplitude-modulated total E-field. Due to the phase difference of the individual E-fields, the total TIS E-field vector also exhibited rotation where the orientations of the two E-fields were not aligned (generally also at the target region). TIS activation thresholds (75–230 V/m) were similar to those of high-frequency stimulation with or without modulation and/or rotation. For E-field dominated by the high-frequency carrier and with minimal amplitude modulation and/or rotation (typically outside the target region), TIS was less effective at activation and more effective at block. Unlike amplitude-modulated high-frequency stimulation, TIS generated conduction block with some orientations and amplitude ratios of individual E-field at very high amplitudes of the total E-field (>1700 V/m).

**Significance:** The complex 3D properties of the TIS E-fields should be accounted for in computational and experimental studies. The mechanisms of suprathreshold cortical TIS appear to involve neural activity block and periodic activation or onset response, consistent with computational studies of peripheral axons. These phenomena occur at E-field strengths too high to be delivered tolerably through scalp electrodes and may inhibit endogenous activity in off-target regions, suggesting limited significance of suprathreshold TIS.

## 1 Introduction

Transcranial electric or magnetic stimulation can modulate the activity of neurons in the brain non-invasively to treat neurological and psychiatric disorders [1–6]. Compared to invasive methods, the electric field (E-field) distribution of transcranial stimulation is more diffuse and shallower due to the large distance between the stimulation sources and the brain and the high conductivity contrast between the tissue layers [7]. For transcranial electrical stimulation (TES), the typical cortical area activated or exposed to significant E-field strength (e.g., more than half the maximum E-field) is on the order of square centimeters [8,9]. For magnetic stimulation (TMS), the fundamental depth-focality trade-off shows that activation of deep targets would result in widespread activation of more superficial brain regions [10,11]. Efforts to improve the spatial characteristics such as focality and depth of conventional stimulation methods rely mostly on improved geometric design of the stimulation devices including electrode montages [12–15] and coil designs [11].

Methods that use the temporal domain and intrinsic neuronal properties were proposed as a means to achieve more focal and/or deep transcranial electrical [16,17] or magnetic [18] stimulation. These methods use multiple stimulation channels, e.g., comprised of separate pairs of scalp electrodes, to generate individual E-fields in the head with different temporal waveforms. Temporal interference stimulation (TIS), originally referred to as interferential current therapy and applied as a neuromuscular treatment between the 1960s and 1990s [19,20], was proposed for non-invasive brain stimulation [16]. In TIS, two high frequency (> 1 kHz) sinusoidal E-fields are applied with a slight frequency difference to create an amplitude-modulated (AM) total E-field through temporal interference. Borrowing terminology from radio technology, neurons are hypothesized to demodulate the low frequency (< 100 Hz) amplitude envelope of the summed E-field waveform via transmembrane current rectification arising from the nonlinear properties of ion channels. Under this assumption, activation would occur mostly in deeper brain regions where the E-field amplitudes of the stimulation channels are similar and therefore the modulation depth (i.e., extent of amplitude change due to the modulation) of the total E-field waveform is large. On the other hand, superficial brain areas would be unaffected, as the mismatch in E-field amplitudes generates a negligible low frequency envelope, and neurons are thought to be insensitive to the large amplitude kHz carrier signal. Without repositioning the electrodes, TIS could steer the site of stimulation by adjusting the relative amplitudes of each channel to shift the region where the E-fields generate the largest amplitude and/or modulation depth of interference [16].

Grossman et al. [16] reported experimental neuronal responses to TIS in the anesthetized mouse brain with current amplitude 2–5 times higher than the threshold for low frequency transcranial alternating current stimulation (tACS), whereas high frequency (kHz) tACS could not evoke responses even at seven times higher current amplitudes. However, the mechanism underlying these effects and the potential application to transcranial stimulation in humans remains unclear. One interpretation is that unmodulated kHz tACS does not significantly affect neurons, putatively due to low-pass filtering at the neuronal membrane, whereas the amplitude modulation exhibited by the summed TIS E-field generates low frequency transmembrane potentials that periodically activate the neurons. However, the neuronal membrane can respond to kHz stimuli since strength–duration time constants of the cortical response to transcranial stimulation are around 150–250 µs [21–23], and membrane filtering does not favor the modulated E-field because the neuronal membrane low-pass filters modulated and unmodulated kHz waveforms to a similar degree [24,25]. Indeed, kHz stimulation can result in transient or sustained firing and, at higher amplitudes, neuronal activity block [25–29]. These responses to kHz stimulation cannot be ignored given that the unmodulated kHz E-field has a stronger absolute amplitude than the low frequency TIS modulation envelope. Even if the effects of the unmodulated kHz stimulus were not detected with specific measurements, this does not necessarily mean that neurons were not significantly impacted. It is possible, for example, that kHz neuronal block suppresses neural responses in regions away from the target [25]. Focality of activation can be achieved in such scenarios, but at the cost of affecting endogenous brain activity in non-target regions.

Several studies explored the mechanisms of TIS, but had limitations in their methods and were not designed to distinguish between the alternative mechanistic explanations outlined above. For example, the Grossman et al. [16] study was in anesthetized mice, and while TIS did not appear to activate off-target brain regions, it is unclear whether it blocked or disrupted endogenous brain activity outside the target. Conduction block by TIS was observed in a model of peripheral nerve stimulation [25]; however, this and other computational studies of suprathreshold TIS focused on activation of isolated axons (fibers of passage) [25,30,31] and did not fully capture effects on cortical neurons. The combination of the two TIS current waveforms results in a three-dimensional (3D) time-varying E-field that can interact in complex ways with 3D cortical neurons [32], but the TIS E-fields and their effect on neurons were analyzed in one dimension (1D), either along the direction of largest modulation depth or in a predetermined direction such as the radial direction of the gray matter or along fiber bundles in the white matter [15,16,25,30,31,33–37]. Some studies explored the modulation of ongoing cortical activity using TIS in the regime of subthreshold field strength of conventional tACS [31,33]; however, they did not provide mechanistic information relevant to deep brain activation by TIS.

Translation of TIS from animal studies to clinical applications poses further challenges. With the same current amplitude, E-fields generated in the human brain are two orders-of-magnitude smaller than those in mice due to differences in head size and cortical geometry. Conventional tACS currents delivered via scalp electrodes are limited to a few mA to be tolerable without anesthesia, generating subthreshold E-fields on the order of only 0.1 to 1 V/m throughout the human brain [15,36,38,39]. Suprathreshold TIS in humans would therefore require much higher current amplitudes. Hence, estimation of TIS E-field thresholds is necessary to provide key information for determining the potential clinical utility of TIS.

The overall organization, goals, and conclusions of this paper are as follows. First, we analyzed the spatiotemporal characteristics of the TIS E-field in 3D, which have been overlooked in the literature (section 2.1). The TIS E-field exhibited some unique properties including frequency jitter and field rotation that may influence neural activation. Second, we simulated activation of individual cortical neurons with TIS and several related tACS modalities (sections 2.2 and 3.1). The E-field thresholds confirmed that suprathreshold TIS delivered with transcranial electrical stimulation would not be feasible in humans because of the high currents that need to be applied to the scalp. The results of related tACS modalities showed that some features of TIS were unique and not reproduced with amplitude modulation and/or field rotation of single-frequency stimulation. Third, we examined whether TIS, and related tACS modalities, could generate conduction block in cortical neurons, as was previously described for peripheral nerve stimulation [25] (sections 2.2 and 3.2). The simulations showed that conduction block of ongoing neural activity can occur and onset response may contribute to or account for the periodic activation observed in TIS in rodents.

## 2. Methods

### 2.1 Characteristics of TIS E-field

Unlike conventional tACS, the E-field from TIS consists of two or more spatial components that have different temporal characteristic. Whereas the E-field of each channel is considered quasistatic, and thus separable into spatial and temporal components (valid up to at least 10 kHz) [40,41], the total TIS E-field is not. Each channels’ E-field can be described as ***E***_*n*_ = ***S***_*n*_**(*s*)** · ***T***_*n*_(*t*), where ***s*** is the macroscopic spatial coordinate vector within the brain, *t* is time, and *n* = 1, 2, … is the channel number. For simplicity, only two-channel TIS is considered and the temporal functions of the tACS channels are normalized: ***T***_*n*_(*t*) = sin(*ω*_*n*_*t* + *α*_*n*_), where *ω*_*n*_ and *α*_*n*_ are their frequency and initial phase. The E-field amplitudes are given according to their spatial distributions |***S***_*n*_|. Specifically for transcranial stimulation, the uniform E-field approximation on the microscopic scale of individual neurons [42] is considered so that ***S***_*n*_ is constant for individual neurons but varies depending on cortical location [43].

### 2.1.1 Aligned E-fields

With *ω*_1_ ≠ *ω*_2_, a 1D AM interferential waveform exists only where the two local E-fields are aligned or anti-aligned (Figure 1A):

**Figure 1.**
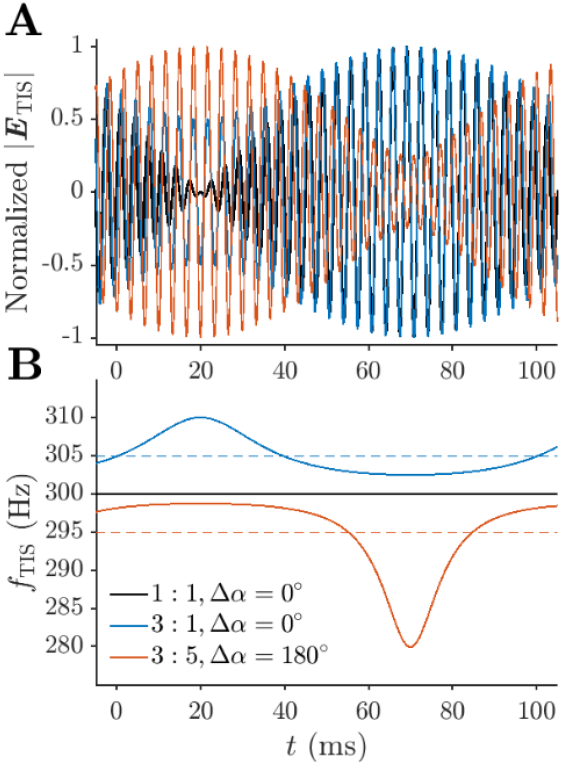
Total TIS E-field consisting of two aligned or anti-aligned E-fields has amplitude-ratio-dependent phase shift and frequency jitter. Characteristic of the total TIS E-field of two aligned (blue and black) or anti-aligned (orange) E-fields with various amplitude ratios. The channels have relatively low frequencies (295 Hz and 305 Hz) for better visualization. **A**. For aligned E-fields (blue and black), the high frequency oscillations of the total E-fields (normalized) are in-phase only at the peaks of their envelopes and shift out of phase at the troughs. For anti-aligned fields (orange), the envelopes themselves are 180° out-of-phase compared to the those of aligned E-fields (blue or black), whereas the carrier could be in-phase at a peak/trough (orange and blue, in-phase at *t ≈* 120 ms) or in between peak and trough (red and black, in-phase at *t ≈* 40 ms). Such phase differences can result in asynchronous neural activation across different cortical locations. **B**. The carrier frequency of the total TIS E-field includes low frequency jitter around the frequency of the E-field with larger amplitude, unlike constant-frequency AM-tACS.

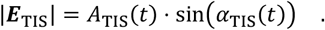

where *A*_TIS_ and *α*_TIS_ are the envelope and instantaneous phase of the oscillation. The envelope

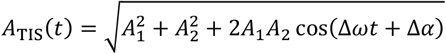

has frequency Δ*ω* = *ω*_2_ − *ω*_1_ and maximum and minimum amplitudes of *A*_1_ + *A*_2_ and |*A*_1_ − *A*_2_|, in which the individual field amplitudes are *A*_1_ = |***S***_1_| and *A*_2_ = |***S***_2_|, and the initial phase difference Δ*α* = *α*_2_ − *α*_1_ is 0° or 180° for TIS (aligned and anti-aligned, respectively). The instantaneous phase is

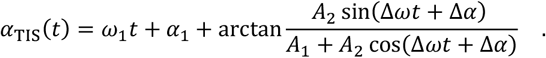

The TIS waveform’s carrier frequency

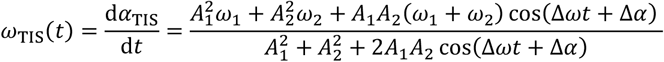

is constant and equals the mean of the two high frequencies only for equal amplitudes. However, for *A*_1_ ≠ *A*_2_, *ω*_TIS_ has low frequency jitter around the frequency of the channel with the larger amplitude (Figure 1B):

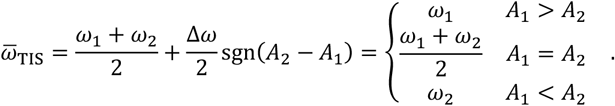

Thus, TIS differs from AM-tACS, which has a constant carrier frequency.

The peak of the low frequency envelope can be out of phase by 180° and the high frequency oscillations have different phase shifts throughout their common low frequency period at different cortical locations, depending on the alignment of the E-fields and their amplitude ratio (Figure 1A).

#### 2.1.2 Non-aligned E-fields

For non-aligned E-fields, the total E-field must be considered in 3D, and not just along the direction of maximum modulation (Figure 2A). In the plane spanned by the two E-field vectors, the total E-field traces out a Lissajous curve bounded by a parallelogram determined by the amplitude, orientation, and phase of the two E-fields (Figure 2B). For E-fields with similar amplitudes and a large angle between them (e.g., 60°-120°), the total E-field alternates between oscillations and rotations at low frequency. The oscillation and rotation correspond to periods in time when the two E-fields are in-phase and out-of-phase and can be characterized by the ratio of the minimum and maximum E-field amplitude during one high frequency cycle.

**Figure 2.**
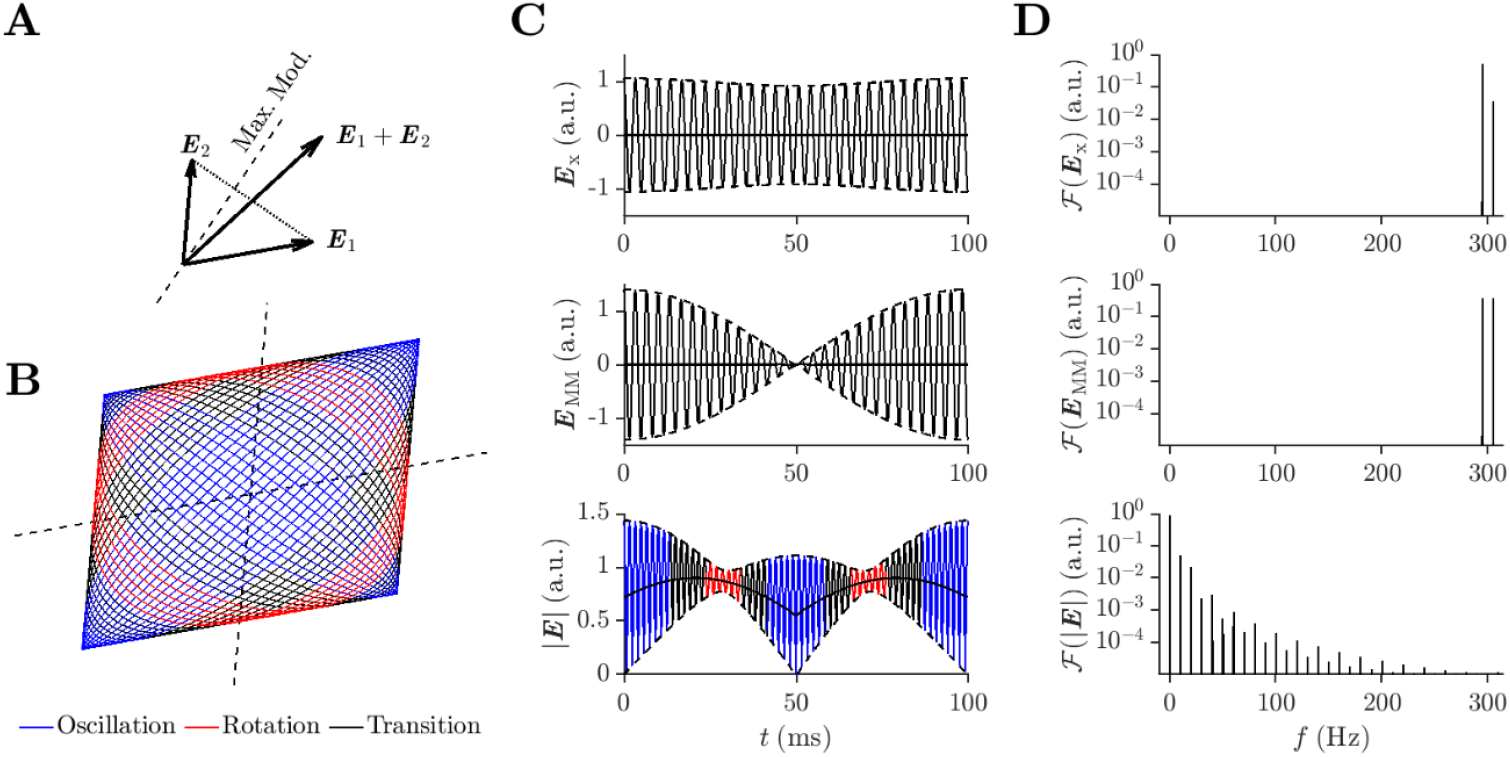
Total TIS E-field consisting of two non-aligned E-fields exhibits field rotation. Characteristics of two non-aligned E-fields with an amplitude ratio of 5:4 and a 75° angle. **A** Considering the E-fields in 3D reveals field rotation previously not considered. The two E-fields of TIS span a plane containing the direction of maximum modulation (Max. Mod., dashed line), which most studies use for analysis of stimulation characteristics, such as amplitude of envelope and modulation depth. **B**. The end point of the total E-field vector traces out a Lissajous curve, which alternates between oscillatory (blue) and rotational (red) periods. Here, oscillation and rotation are (arbitrarily) defined as the E-field magnitude having a ratio of minimum versus maximum smaller than 1/3 or larger than 2/3 during one high frequency cycle, respectively. **C**. The total E-field projected along any direction, for example the *x*-axis (***E***_*x*_, top row) or the direction of maximum modulation (***E***_MM_, middle row), has a low frequency envelope (dashed lines). The oscillations, however, average out to zero (thick line) and therefore the waveform does not contain low frequency energy. In contrast, the E-field magnitude (bottom row) has low frequency content that could be a driving factor of TIS. **D**. The corresponding spectrum of the E-fields in C. ℱ: Fourier transform. a.u.: arbitrary units.

Unlike any 1D E-field projection that does not contain low-frequency content [25], the amplitude of the rotating E-field does contain low-frequency power (Figure 2C and 2D) and field rotation could contribute to stimulation by TIS. For example, multi-electrode tACS with phase-shift generates global traveling waves and local rotation for the E-field [44], and rotational field TMS, applied using two perpendicular figure-of-eight coils driven with the same current waveforms with a 90° phase shift, exhibited lower thresholds and stronger evoked responses [45–47]. The rotational E-field is hypothesized to sweep across the neural population and activate neurons with different directional sensitivity [45,47].

Given the large difference between the volumes of brain where the two TIS E-fields are (approximately) aligned versus not aligned, 1D parameters such as modulation depth, carrier frequency, and envelope do not encompass the full spectrum of TIS scenarios. Capturing the spatial variations in E-field dynamics within individual neurons in computational simulations requires neuronal models with realistic morphology instead of point neuron models. For a given amplitude ratio between the stimulation channels, only the ratio of the E-field amplitude can be specified for any neuron under the quasi-uniform approximation, and no single modulation depth can be defined because the waveform of the E-fields projected along the neural cables varies for different neuronal compartments.

### 2.2 Neuronal response to TIS and other forms of tACS

#### 2.2.1 Stimulation waveforms

Three conventional (static) and three non-static tACS methods were studied (Figure 3). The conventional methods comprising a single static E-field included 10 Hz low frequency stimulation (LFS), 2 kHz high frequency stimulation (HFS), and 2 kHz AM-HFS with 10 Hz modulation. Non-static methods included TIS (two E-fields with 2 kHz and 2.01 kHz frequency, respectively), rotational field HFS (RFS, two E-fields with 2 kHz frequency and a 90° phase shift) and AM-RFS (RFS with 10 Hz-modulated E-fields), which were all simulated with E-field amplitude ratios of 1:1, 2:1, 4:1. RFS was studied to examine how the field rotation alone affects tACS, and AM- RFS was included to examine how non-interference-generated AM compares with AM of TIS. The waveforms had a duration of 4 s and, to reduce onset response, 0.5 s and 0.25 s linear ramp-up and ramp-down at the beginning and end, respectively.

**Figure 3.**
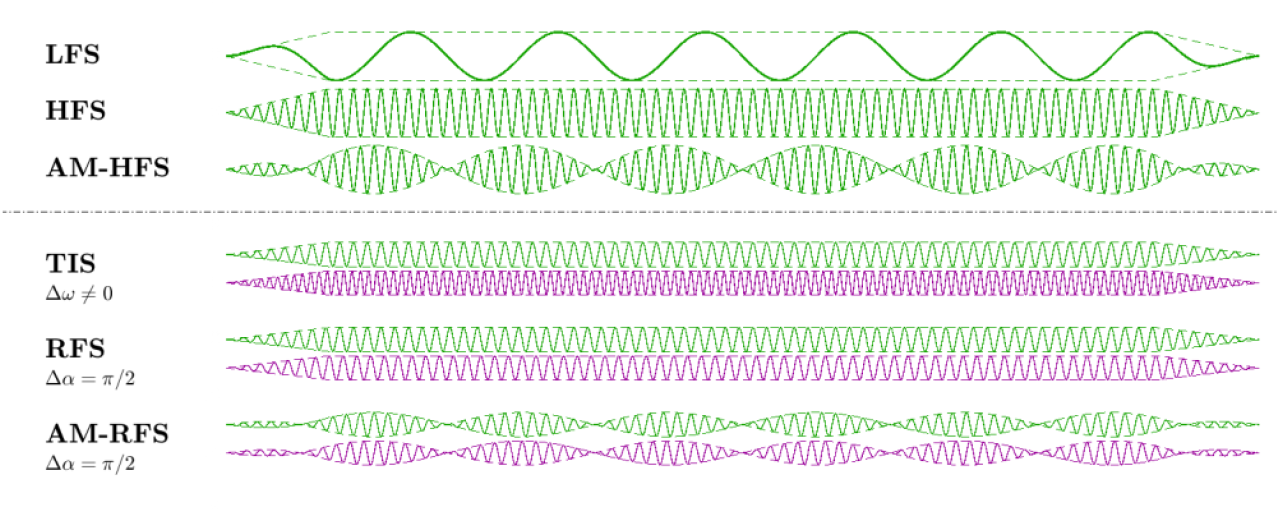
Stimulation waveforms of various tACS modalities. Illustration of the waveforms of low frequency stimulation (LFS), high frequency stimulation (HFS), amplitude-modulated HFS (AM-HFS), temporal interference stimulation (TIS), rotational field HFS (RFS), and amplitude-modulated RFS (AM-RFS). The waveforms of the conventional methods and the first E-field of non-static stimulation methods are shown in green, and the waveform of the second E-field of non-static stimulation methods are shown in purple. The envelope of all waveforms is shown with dashed lines and has a ramp-up and ramp-down period. The relative frequencies are for visualization and not to scale with the actual stimulation.

#### 2.2.2 Models of realistic neurons

To capture fully the effect of TIS, RFS, and AM-RFS on 3D cortical neurons, we used biophysically realistic neuron models from the Blue Brain project [48,49] that were previously adapted and validated for stimulation with extracellular E-fields [32,43]. The models were implemented in the NEURON v7.4 [50] and included cell-type specific 3D dendritic and axonal morphologies reconstructed from juvenile rat brain slices and up to 13 Hodgkin-Huxley-like ion channel types. Key modifications included myelinating the axons, scaling ion channel kinetics to body temperature, and assigning ion channel properties to the entire axon arbor [43]. Layer 5 (L5) pyramidal cells (PC, Figure 4A) were simulated as they have the largest size and thus lowest thresholds in response to extracellular stimulation [32,43]. The main axon’s terminal was disabled from direct activation by extracellular stimulation to account for the truncation of the main axons in the slicing process and the artificial generation of low threshold sites [32]. To capture better the response to kHz stimulation and match more recent studies of axonal excitability [51,52], the kinetics of the sodium channels were accelerated by uniformly scaling the time constants of the gating variables of the fast sodium channel (*m*^3^*h*) by 0.4 for both the activation (*m*) and inactivation (*h*) gates. The overall effect was a faster firing rate in response to intracellular stimulation, and a decrease in thresholds to kHz stimulation (in the range of −20% to −12.5%) with limited effects on LFS (Figure S1). We conducted simulations of all stimulation parameters with one representative L5 PC model and confirmed that the findings are generalizable across cell morphologies using simulations of a subset of parameters in four additional model clones (Figure S2). We also tested the thresholds of a L2/3 PC model and a L4 large basket cell model (Figure S2), but we did not carry out more extensive characterization of these cell types since their thresholds were significantly higher than those of the L5 PCs for both low and high frequency stimulation.

**Figure 4.**
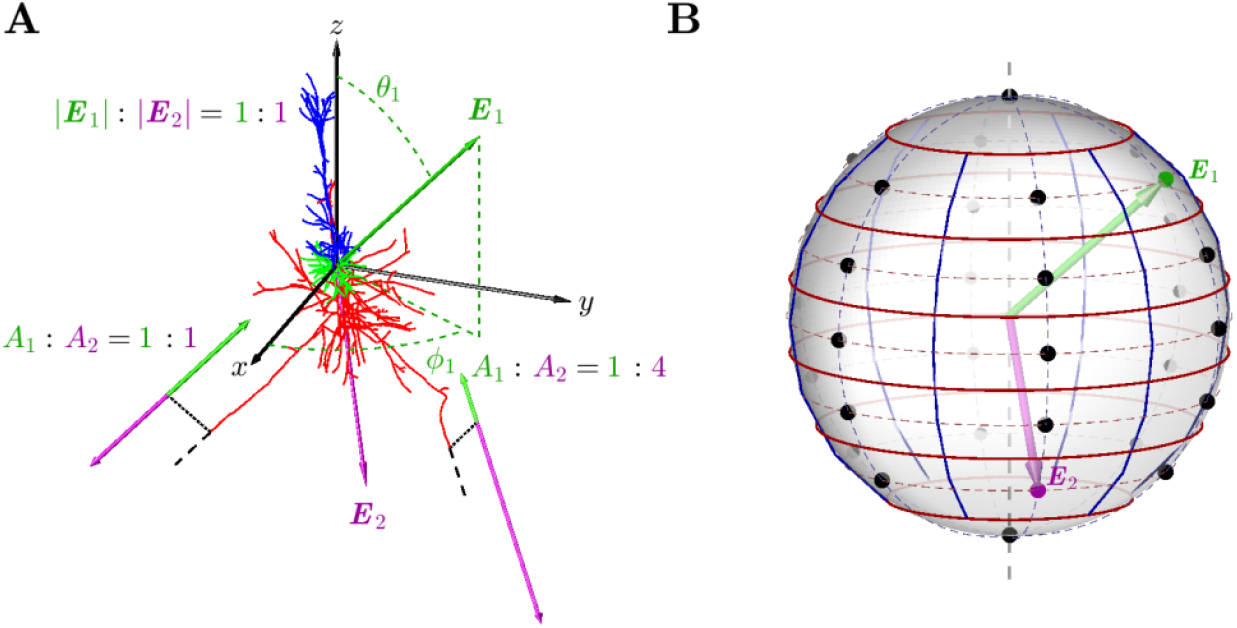
Model neuron and definitions of E-field orientations in 3D. A. L5 PC morphology and the spherical coordinate system for E-field orientation. The origin is located at the soma with the *z*-axis aligned along the somatodendritic axis. The E-fields are defined by their polar and azimuthal angles, *θ* and *ϕ*, shown for the first E-field (green) as example. The E-fields have different effective amplitudes (*A*_*1*_, *A*_*2*_) at different locations along the neural cable, as shown at two terminals in the axonal arbor, with the black dashed lines showing the orientation of the axon at the terminals. Color legend for neuron: apical dendrite in blue, basal dendrite in green, and axon in red. **B**. Sampling of 32 E-field orientations for the 3D neuron model represented by dots on a spherical surface, with black and grey dots indicating the front and back of the sphere. The polar axis is shown as a vertical dashed line.

#### 2.2.3 Simulation setup

Following the uniform E-field approximation discussed in Section 2.1, the orientations of the E-fields were defined in a spherical coordinate system centered at the soma and with the polar direction (*z*-axis) aligned with the somatodendritic axis (Figure 4A) [43]. These orientations can represent any possible cortical neuron locations for any E-field distribution, regardless of the electrode montage. Given the alignment of the somatodendritic axis with the cortical column, the polar angle reflects the orientation of the E-field relative to the cortical geometry and more significantly influences activation thresholds, whereas the azimuthal rotation of neuron models within the cortex are random [32]. To achieve a uniform representation on the spherical surface, the 32 E-field orientations were taken from a grid of seven non-uniformly sampled polar angles and six uniformly sampled azimuthal angles (Figure 4B). For stimulation methods comprising two E-fields, both fields were sampled with the 32 orientations, yielding a total of 1024 (32 × 32) orientation combinations for each parameter setting (i.e., frequency, amplitude modulation, phase shift, and amplitude ratio). NEURON’s extracellular mechanism [43,50] was used for coupling the E-fields to the models, with the extracellular potential *V*_e_ for each compartment *C* calculated as

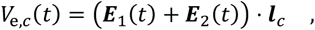

where ***l***_*C*_ is the displacement vector specifying the compartment location relative to the soma. The simulations were set up in MATLAB (versions 2019a and 2020a, The Mathworks, Natick, MA, USA) and performed in NEURON with a time step of 10 μs, which was sufficient to capture the approximately 0.5 ms period of the kHz stimulation and the response of the fast sodium channels with fastest time constant of ∼ 40 μs.

The threshold for activation was defined as a response with a firing rate of at least 9.5 Hz at the soma, which is 95% of the 10 Hz frequency of LFS, the difference frequency of TIS, or the frequency of the amplitude modulation (Figure 5A). Bursting was defined as spiking events with less than 15 ms inter-spike intervals within a 50 ms duration [16]; if bursting occurred, the bursts were counted instead of individual spikes to calculate the firing rate in the threshold search. Only spikes within the steady state period of the stimulation were counted, and onset or offset response during the ramp-up and ramp-down time was not considered for threshold determination. Therefore, a suprathreshold response should have at least 31 spikes or bursts during the steady state stimulation period. Setting the threshold firing rate to a slightly lower value allowed the activity in model neurons to mimic experimental recordings, in which some trials did not result in firing during every cycle of the LFS or TIS envelope [16]. Threshold amplitude was determined by a binary search to 1% accuracy and, for non-static stimulation methods, was defined as the maximum amplitude of the total E-field. To determine whether action potentials were initiated in the soma or axons, we compared the spike timing at the soma, axon initial segment, first node of the axon (directly connected to the initial segment), and terminal of the main axon.

**Figure 5.**
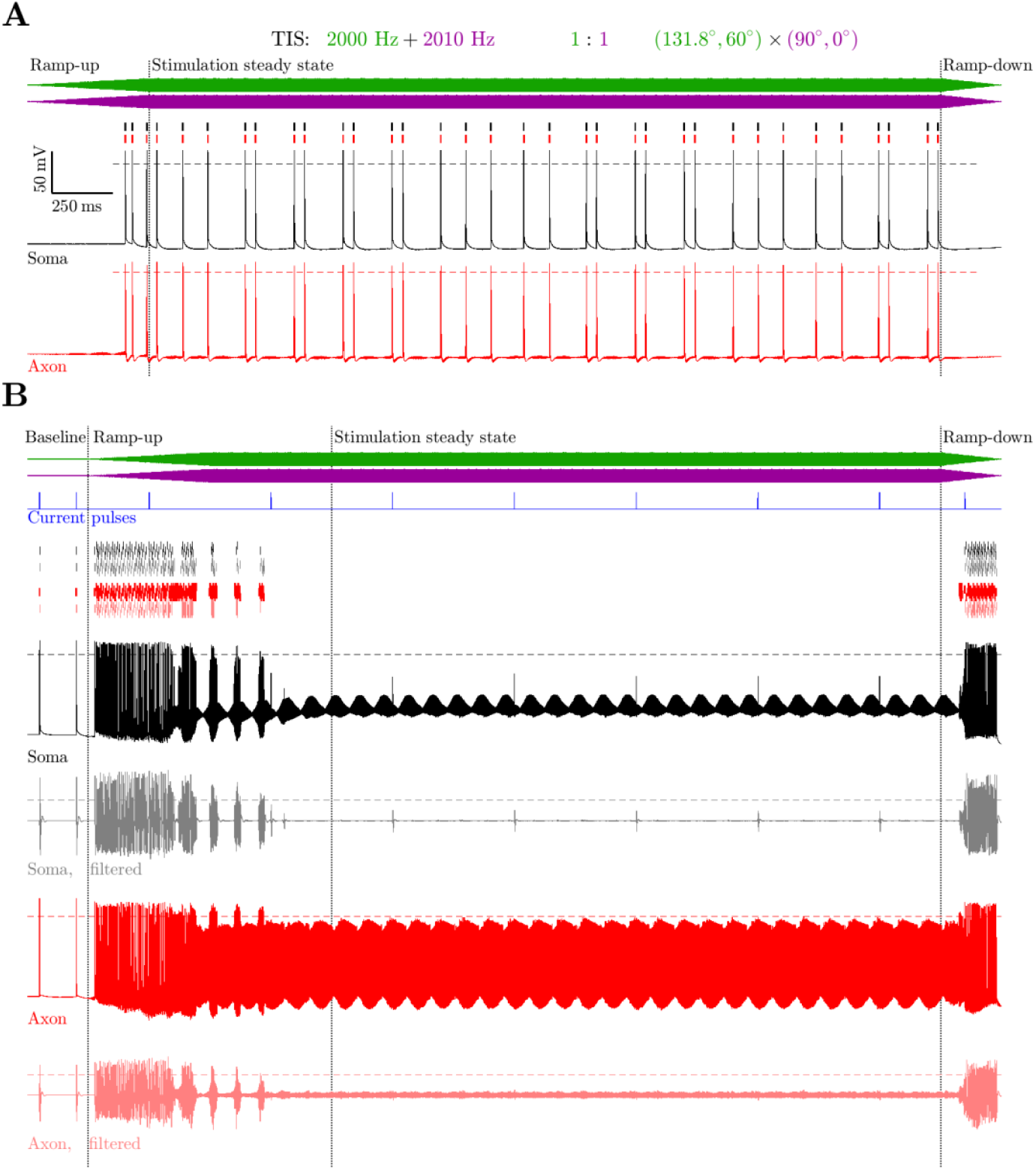
Neural response and neural recording processing. **A**. Example response of L5 PC firing in response to TIS at threshold. The stimulation waveforms of the two E-fields are shown on top, followed by raster plots of firing and transmembrane potentials for the soma (black) and an axon node (red). Individual spikes or the first spike within a burst are aligned horizontally, with additional spikes within a burst offset vertically. The dotted vertical lines mark the start and end of the stimulation steady state period between 0.5 s and 3.75 s, and the horizontal dashed line marks the spike thresholds. **B**. Example response of L5 PC conduction block in response to TIS at high amplitude. Suprathreshold intracellular current pulses (blue) were applied to the soma and evoked action potentials at baseline. However, they did not evoke action potentials during the stimulation steady state, either in the raw (black) or filtered (grey) membrane potential. The membrane potential deflection in the soma generated by the test current pulse did not generate action potential in the axon (red traces). The dotted vertical lines mark the baseline period ending at 0.25 s and start and end of the stimulation’s steady state period between 1 s and 3.75 s.

Thresholds for conduction block were determined to examine whether the non-static stimulation methods generated high frequency block in model cortical neurons, as was observed in models of peripheral axons [25]. Suprathreshold test current pulses were injected into the soma to examine transmission of action potentials. The start of waveforms was therefore delayed by 0.25 s to allow test pulses to be applied during a baseline period. To accommodate the strong onset response, the steady state was considered to start 0.25 s after the amplitude ramp-up ended. Therefore, the period for which action potentials were considered for conduction block was 2.75 s, starting 1 s after the stimulus onset. The threshold search was terminated and “no block” was determined for a particular set of stimulation parameters when the maximum tested amplitude of the (first) E-field reached 6000 V/m and action potentials were still occurring during the steady state stimulation period. This upper limit was chosen as an arbitrarily high E-field value well above the stimulation threshold and about three times larger than the lowest block threshold observed in the simulations. To match the filtering that was applied to patch clamp recordings of cortical and hippocampal neurons [16], the membrane potential was filtered using a 5^th^ order band stop filter (1 kHz to 15 kHz) and a 3^rd^ order high pass filter (100 Hz). This removed the stimulation “artifact” due to the high stimulation amplitude directly driving the neural membrane above the 0 mV threshold, revealing the underlying action potentials (Figure 5B). Firing patterns were compared for both the raw and filtered transmembrane potential, with a 25 mV threshold used for the latter.

## 3 Results

### 3.1 Firing pattern and threshold of activation

For all tACS methods and across all E-field orientations or combinations of orientations, spiking events occurred earliest in the first node of the axon. As axon terminals had the lowest thresholds to extracellular stimulation [43,53], the responses were all initiated in the axonal arbor and propagated antidromically to the soma and re- orthodromically through the main axon [32,43].

At threshold, the firing patterns evoked by all stimulation modalities showed a response before the E-field oscillations reached steady state amplitude (Figure 6), and this was not eliminated by the ramp-up of the E-field magnitude(s). For LFS, AM-HFS, and AM-RFS, the responses during stimulation steady state were single spikes at regular intervals corresponding to the frequency of the stimulation or amplitude modulation. For HFS, TIS, and RFS, the bursts of activity before stimulation reached steady state appeared to be a transient onset response. During steady state stimulation, the HFS and RFS responses consisted of short patterns of two spikes that repeated regularly but did not satisfy the definition of bursts. The TIS responses exhibited regular firing at the difference frequency for most combinations of E-field orientation, and changing the amplitude ratio had negligible effect on the firing pattern. For some combinations of orientation, the firing pattern was irregular and exhibited bursts consisting exclusively of double spikes. For TIS with a 1:1 E-field amplitude ratio, bursting occurred in 229 of the 1024 combinations of orientation of the E-fields. In combinations where bursts occurred, bursting accounted for 3.2 ± 0.1% of firing events (mean ± standard deviation), whereas among firing events across all combinations of orientation, only 0.7% were bursts. Increasing the E-field amplitude ratio to 2:1 and 4:1 reduced both the number of combinations evoking bursting and the number of bursts (181, 3.2 ± 0.1%, and 0.6%, and 78, 3.3 ± 0.1%, and 0.2%, respectively). While only TIS evoked bursting at threshold, the neuron model could respond with bursting to suprathreshold stimulation amplitudes for all stimulation modalities (Figure S3).

**Figure 6.**
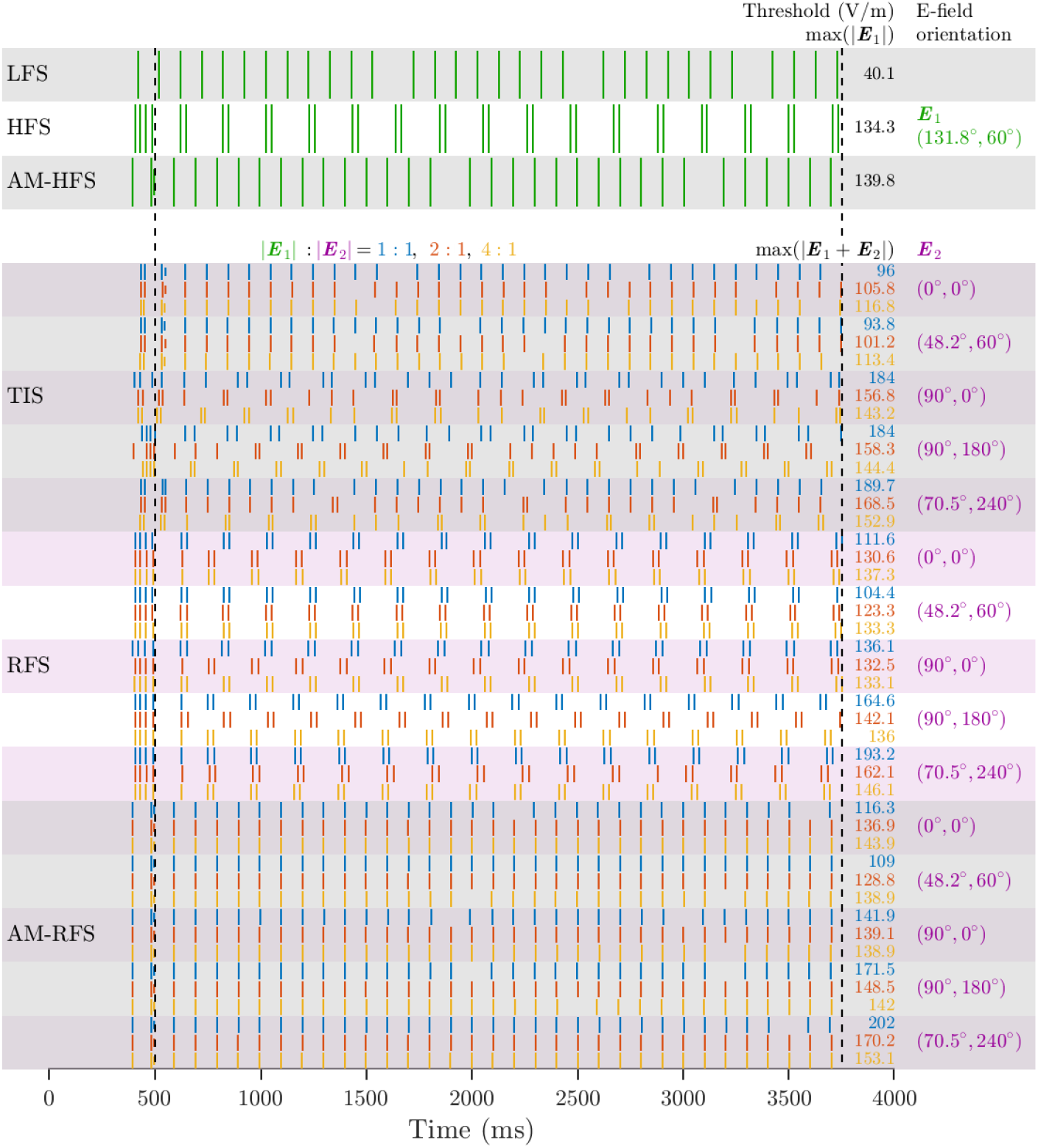
Representative responses of L5 PC at activation threshold. Raster plot of somatic firing shows periodic activation by all tACS modalities, with HFS, TIS, and RFS generating a pattern of two action potentials in rapid succession. The same orientation was chosen for the E-field of LFS, HFS, and AM-HFS as well as for the first E-field of TIS, RFS, and AM-RFS. The same five orientations were chosen for the second E-field of TIS, RFS, and AM-RFS, with results for all 3 amplitude ratios of the two E-fields shown (from top to bottom: 1:1, 2:1, 4:1 in blue, red, and yellow, respectively). The dotted vertical lines mark the start and end of the stimulation steady state period between 0.5 s and 3.75 s.

Activation thresholds were significantly lower for LFS (range 16.9 V/m to 47.4 V/m) compared to all kHz tACS modalities (Figures 7 and S4**-**S8), as expected due to the low-pass filtering by the neuronal membrane [24,25,29]. The kHz tACS thresholds all fell within a similar range (75 V/m to 230 V/m) despite some variation in their distributions for different E-field orientations. The threshold for a stimulation modality corresponded to the lower range of the distributions (∼ 75 V/m to 100 V/m), because for the large and dense population of cortical neurons represented by the randomly-rotated model cells, there were always some neurons that exhibited the lowest threshold. The thresholds were similar to TMS thresholds [32,54], despite the latter using sub-ms pulsed stimulation, indicating that the continuous aspect of the waveforms did not contribute critically to the threshold. Thresholds varied with the E-field orientations (Figures S5-S8). Due to the oscillation of the E-fields, the threshold maps were symmetric for points corresponding to E-fields of opposite orientations. The lowest threshold orientation for LFS was aligned with an axon collateral, whereas the lowest threshold for kHz tACS was aligned with the somatodendritic axis. For TIS, RFS, and AM-RFS, the thresholds mostly increased with the E-field amplitude ratio (Figure S4). The change was less prominent for TIS due to the response being dominated by the first E-field, and this also resulted in clustering of thresholds for unequal E-field amplitudes (Figure 7).

**Figure 7.**
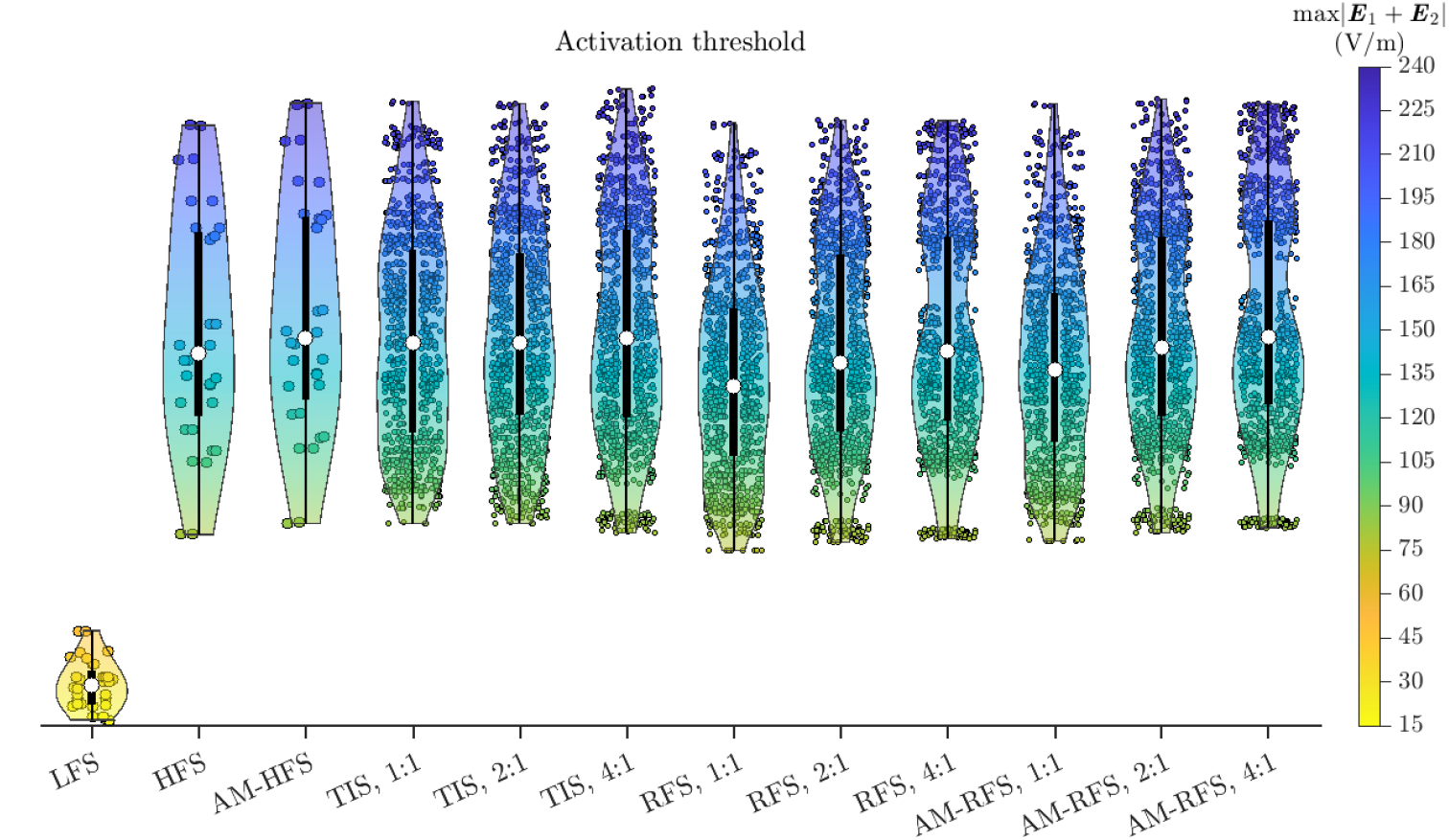
Activation thresholds are similar for all kHz stimulation modalities. Thresholds for action potential generation for all 32 orientations (LFS, HFS, and AM-HFS) and 1024 orientation combinations (TIS, RFS, and AM-RFS) of E-field(s). Each violin plot shows the individual data points as circles (with jitter fixed for the E-field orientations and orientation combinations), the estimated density of distribution as a shaded area, and several statistics as box-and-whisker, with the white circle representing the median, the box covering the first to third quartiles, and the whiskers extending to the upper and lower adjacent values that are 1.5 times the interquartile range above and below the quartiles.

### 3.2 Firing pattern and threshold of conduction block

At sufficiently high E-field amplitudes, all kHz tACS modalities evoked a strong onset response during ramp up and an offset response during ramp down (Figures 8, 9, and S9). For the majority of E-field orientations of HFS, TIS, and RFS, conduction block could occur during the stimulation steady state (Figures 10 and S10-S14). For all AM-HFS and AM-RFS cases, periodic activation at the frequency of amplitude modulation occurred for all suprathreshold E-field amplitudes. At low stimulation amplitudes, the activation coincided with the maxima of the AM envelope. At high amplitudes, however, the activation shifted to the minima of the envelope, indicating the activation was an onset/offset response separated by conduction block during periods of high E-field amplitude (Figure S14A and S14D–E). Similarly, when sustained block was not observed for TIS, the bursts of firing were shifted by half the modulation period relative to the burst activation at lower field strengths, and the neural membrane was blocked between the onset and offset responses (Figures 9C).

**Figure 8.**
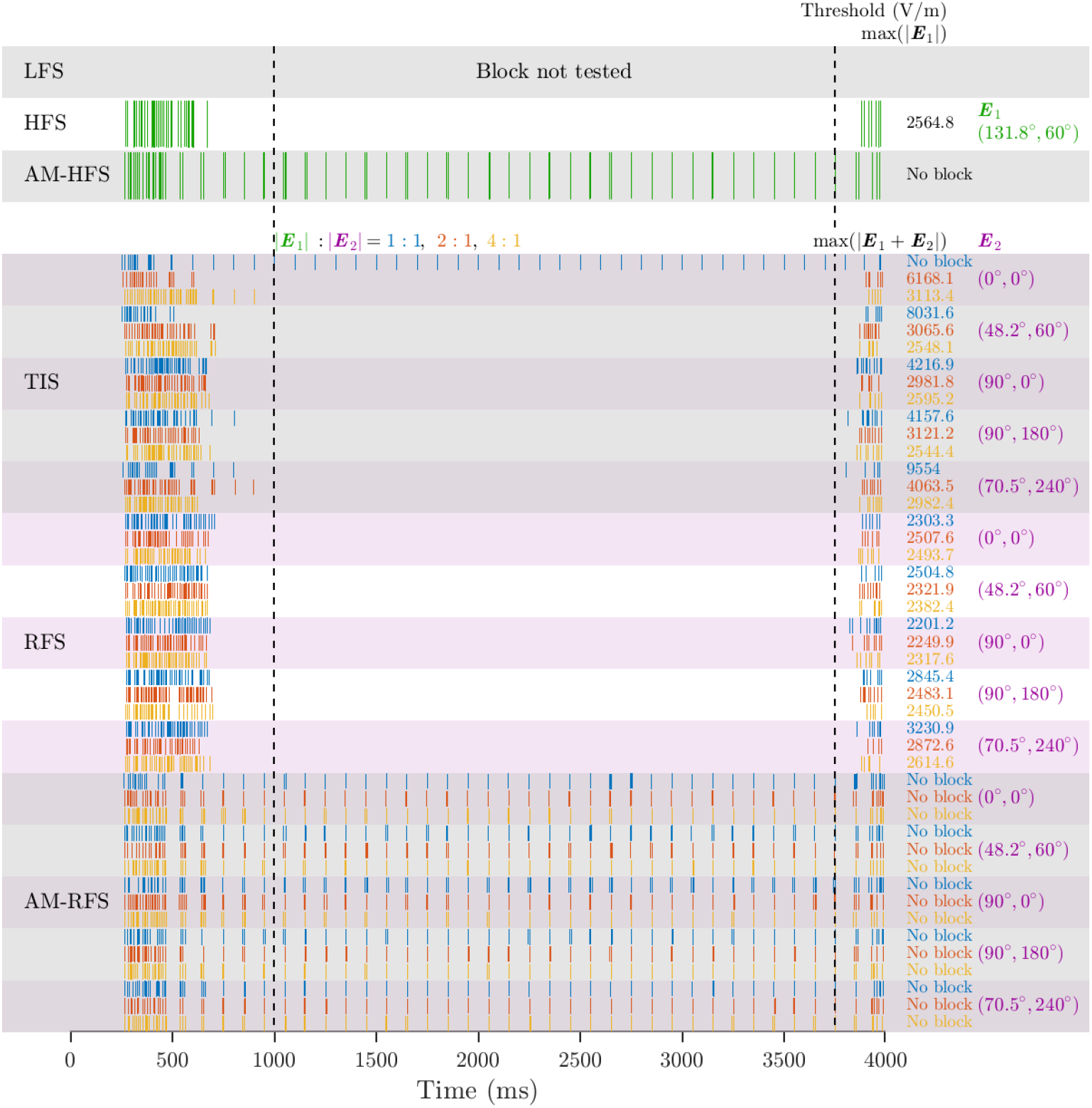
Representative responses of L5 PC at conduction block thresholds or maximum amplitude tested. Raster plot of somatic firing shows that conduction block occurred in response to HFS, RFS, and some TIS orientation combinations, but not for AM-HFS or AM-RFS. The spikes of the filtered transmembrane potential of the soma are shown for the same E-field orientations as in Figure 6. Results for spikes of the raw transmembrane potential are shown in Figure S8.

**Figure 9.**
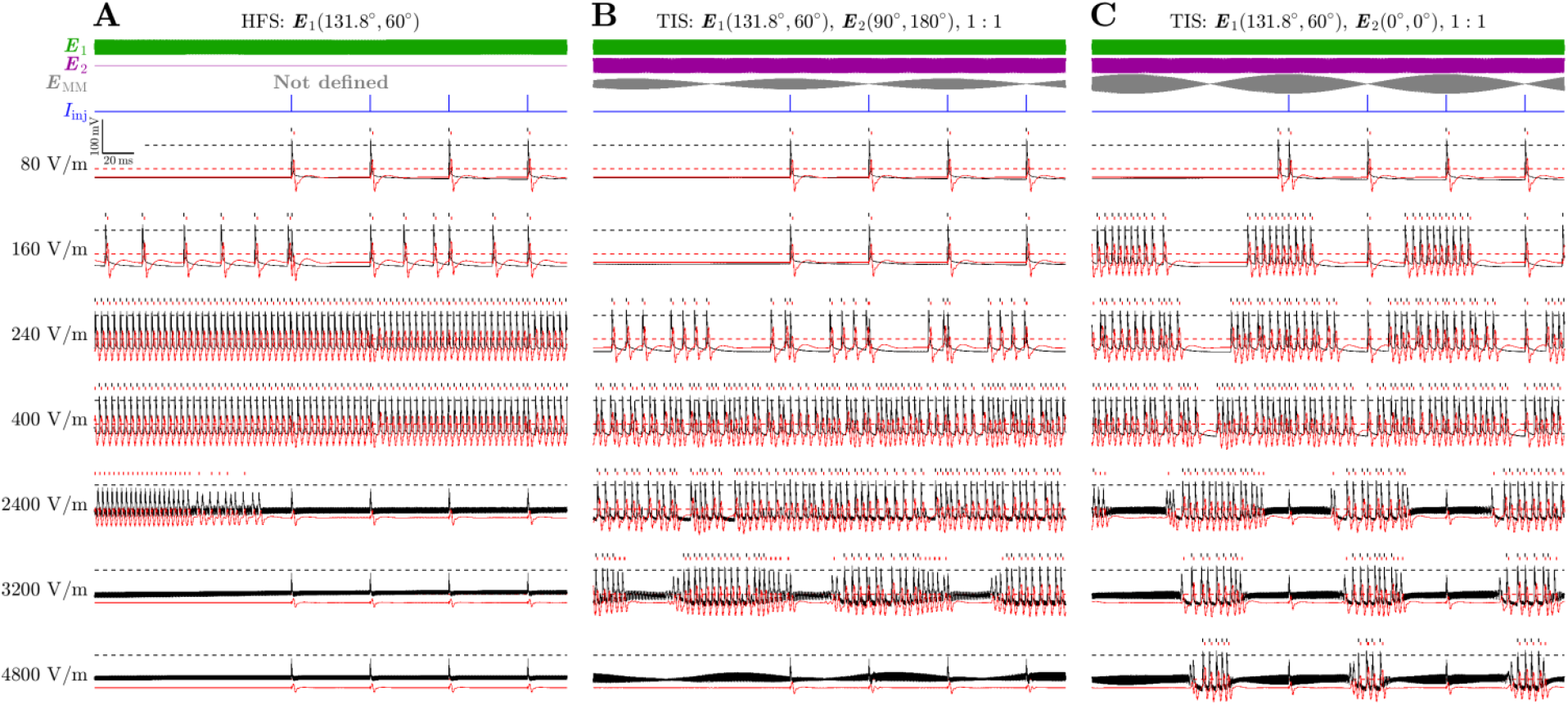
Representative neuron firing evoked by HFS and TIS for a range of E-field amplitudes demonstrates conduction block and shift in activation for TIS with higher amplitudes. The raw (black) and filtered (red) transmembrane potential at the soma with action potential timing marked with vertical lines for seven different E-field amplitudes given on the left as either max(|***E***_1_|) for HFS or max(|***E***_1_+***E***_2_|) for TIS. The dashed lines show the action potential threshold of 0 and 25 mV for the raw and filtered transmembrane potentials, respectively. Shown on top are the individual E-field waveforms (green and purple), total E-field along the direction of maximum modulation (***E***_MM_, gray), and timing of suprathreshold current injection test pulse (*I*_inj_, blue). The orientation of the (first) E-field is the same as in Figures 6 and 8, and two orientations of the second E-field are shown. **A**. HFS, which is equivalent to TIS with 1:0 amplitude ratio, evoked sustained firing with increased E-field amplitude and resulted in conduction block at high stimulation amplitudes. **B–C**. TIS evoked periodic firing at lower amplitudes, which became sustained firing similar to HFS at higher amplitudes, and then became periodic again at even higher amplitudes as the amplitude modulation of the total TIS E-field resulted in onset and offset firing. Depending on the E-field orientations and corresponding amplitude modulation of the total E-field, the onset/offset firing may cease (B) or persist (C) at higher E-field amplitudes, but the transmembrane potential reached a blocked state and did not respond to stimulation. Results for AM-HFS, RFS, and AM-RFS are shown in Figure S14.

**Figure 10.**
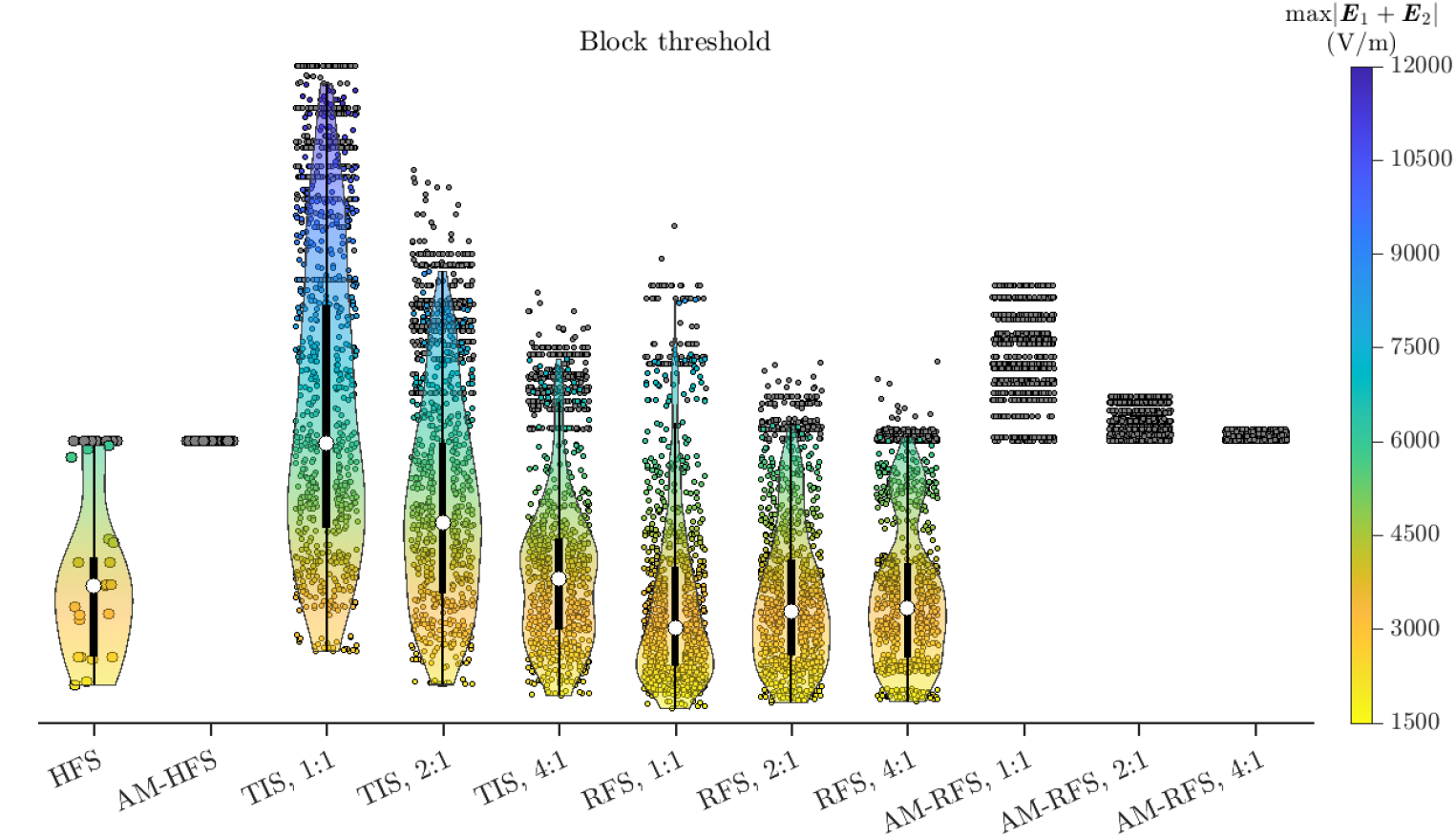
TIS can generate conduction block at overall higher thresholds compared to HFS and RFS, whereas AM methods do not cause block. Thresholds for all 32 orientations (HFS and AM-HFS) and 1024 orientation combinations (TIS, RFS, and AM-RFS) of E-field, in a format similar to Figure 7. Orientations (combinations) that did not achieve block with |***E***_1_| ≤ 6000 V/m are shown as grey circles.

Block thresholds had a wide distribution (Figure 10) and were strongly dependent on E-field orientation (Figures S10-S13). Block thresholds for TIS decreased with increasing E-field amplitude ratio, and, except for 4:1, were higher compared to those of HFS and RFS. For TIS with equal E-field amplitudes, neurons were more likely to be exposed to a total E-field with stronger low-frequency modulation (Figure 4A), which generated periodic onset and offset responses interrupting the block. On the other hand, for unequal E-field amplitudes, the modulation was weaker and therefore could achieve block at lower thresholds. Block thresholds for RFS increased with amplitude ratio and were similar to those of HFS. For RFS, equal E-field amplitude exposed more axon terminals to higher aligned E-fields at the kHz frequency, resulting in block at lower thresholds. In contrast, some axon terminals were aligned with the weaker E-fields of the unequal E-fields, resulting in higher block thresholds.

## 4. Discussion

### 4.1 Characteristics of TIS

We quantified the activation and block of model cortical neurons by single channel tACS and by methods for modifying the spatial selectivity of tACS using two high-frequency currents to generate E-field temporal interference with two closely spaced frequencies or E-field rotation through relative phase shift without or with amplitude modulation. TIS evoked periodic activation of the model neurons at the difference frequency of the two stimulation currents, similar to prior results [16]. However, unlike the experimental results, TIS thresholds were comparable to those of HFS, as well as those of RFS or their AM versions. The E-field amplitudes to achieve activation of model neurons with all forms of kHz tACS (75 to 230 V/m) were much higher than the range for LFS (16.9 to 47.4 V/m). At markedly higher amplitudes (> 1700 V/m), neuronal responses and endogenous activity could be blocked by non-modulated kHz tACS, including TIS. While TIS did generate amplitude modulation of the total E-field projected locally along the neural elements, the depth of modulation and magnitude of the modulated waveform were highly dependent on the amplitude ratio of the two E-fields as well as their orientations with respect to the neural element. Therefore, the E-field projected on the neural elements has smaller magnitude and shallower modulation compared to the maximum values determined by the TIS E-fields alone. Consequently, TIS generates conduction block similarly to regular kHz stimulation without modulation, whereas AM-tACS with maximum modulation depth did not generate block for any E-field direction.

Our study related neuronal responses to the strength and temporal characteristics of the local E-field. To link the transcranial stimulus current with the E-field, Rampersad et al. [36] simulated the E-field in a mouse brain with a model representing the conditions of the original study [16]. They reported a TIS E-field maximum of 383 V/m, higher than the threshold E-field values in our study. However, their E-field was defined for the amplitude of the envelope along the orientation of maximum modulation [16,36]

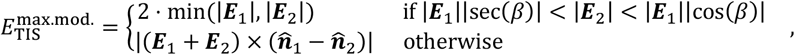

where *β* ∈ [0, π] is the angle between the two E-fields and 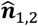 are unit vectors along the E-fields. The E-field along the maximum modulation orientation does not reflect the maximum E-field, max|***E***_TIS_(*t*)|, over an entire low frequency period of TIS and, in general, is smaller than the latter (Figure 1B). A metric comparable to our reported thresholds is the maximum E-field amplitude of conventional two-channel tACS with the same combination of E-field orientations and amplitude ratio, which was 38,000 V/m [36]. This value is much higher than our activation thresholds and an order of magnitude higher than the conduction block thresholds. The model by Rampersad et al. did not include the skin, and such a high E-field amplitude compared to other studies [38] may be attributed to the absence of current shunting by conductive soft tissues [7]. Nevertheless, even if Rampersad et al. overestimated the E-field by tenfold, our simulation results indicate that TIS applied epicranially to mice could achieve conduction block in cortical neurons, similar to previous computational studies that examined the effects of TIS on peripheral axons [25]. Therefore, the periodic activation observed *in vivo* in mouse experiments may reflect onset responses of nerve block in the target region where the two E-fields have similar amplitudes. The block was released periodically in the target region due to amplitude modulation of the TIS E-field, whereas the surrounding area was completely blocked due to much smaller modulation depth.

In humans, however, tolerable tACS currents for scalp stimulation without anesthesia are only a few mA and generate E-fields in the brain of only 0.1 V/m to 1 V/m [8,39]. While these E-field amplitudes are sufficient to modulate neural activity [17,55–57], they are orders of magnitude smaller than both activation and block thresholds of TIS. Our modeling results indicate that suprathreshold TIS would require E-fields in the range from 150 V/m to 800 V/m, corresponding to scalp electrode current of hundreds of mA to over 1 A, which is comparable to or exceeds that in electroconvulsive therapy (ECT) [58,59]. Therefore, the generation of suprathreshold TIS effects would be intolerable without anesthesia, could result in heating at the electrode–scalp interface, and may induce a seizure with continuous application. Nevertheless, subthreshold TIS [31,33], which involves very different mechanisms, may still merit investigation due to the unique properties of the TIS E-field. Likewise, TIS could be of interest in suprathreshold applications under anesthesia in the context of ECT or intensive nonconvulsive stimulation [60] as well as for invasive stimulation [61].

Our results revealed complexities of the E-fields produced by TIS and RFS that have implications for the design and interpretation of modeling and experimental studies. The activation and block thresholds for the E-fields of these non-static stimulation methods varied not only with their amplitude ratio and angle but also with their orientations relative to the neuron morphology. For example, ***E***_1_ and ***E***_2_ with polar angles (*θ*_1_ and *θ*_2_) and co- varying azimuths (*φ*_1_*−φ*_2_ = constant) span a fixed angle, but the thresholds vary substantially (Figure S6-S8 and S12-S13). Therefore, the E-fields obtained along specific directions in computational models, whether for maximum modulation [15] or in a pre-determined orientation [16,36,37], cannot predict the stimulation effects of TIS, and optimization using calculation of the E-field alone [35,62] is unlikely to produce the same results as when considering the response of the neurons to those E-fields.

Further, TIS has unique features compared to the other kHz stimulation methods that were analyzed. TIS creates amplitude modulation in regions where the two E-field amplitudes are similar, which generally correspond to the stimulation target. The slight differences between the TIS waveforms for aligned E-fields compared to that of a single channel kHz AM-tACS could have non-negligible effects on neural activation. TIS also creates a rotational E-field where the two E-field orientations are not directly aligned, and this generally includes the target region as the two pairs of electrodes are placed on opposite sides of the head. Design or optimization of electrode montages that exploit non-static fields [15,62] could benefit from including field rotation. In contrast, the E-field of conventional AM-tACS contains no rotation and would only engage a subset of the neural targets recruited by TIS, despite being able to achieve larger modulation depths for a given orientation [15]. The rotation of TIS is emergent due to the varying phase difference during the low frequency cycle, whereas the magnitude of rotation of RFS is sensitive to the phase difference between the waveforms applied at the two pairs of electrodes. The total TIS E-field is mostly oscillatory with minimal rotation and amplitude modulation when one E-field has much larger amplitude than the other, which typically occurs in regions outside the target. This results in similar activation thresholds to HFS and could result in conduction block at high field amplitudes, where higher amplitude ratios would confer lower block thresholds than in the target region. Of the methods capable of producing conduction block, TIS also has the highest block thresholds. AM-HFS and AM-RFS cannot achieve conduction block, whereas RFS achieves block at lower thresholds for equal E-field amplitudes compared to unequal amplitudes. On the network level, the asynchronous variation of the TIS E-field over space should be considered [44]. Therefore, experiments such as brain slice studies should consider replicating the features of the TIS E-field with either two stimulation channels or a simultaneously amplitude- and frequency-modulated E-field, rather than substituting it with a single-frequency AM E-field [33].

### 4.2 Filtering of transmembrane potentials

We analyzed the neuronal responses based on the raw transmembrane potential recordings to determine activation thresholds. At threshold E-field amplitudes, the kHz oscillations of transmembrane potential had limited amplitudes (5 mV to 15 mV) throughout the cell and did not mask the action potentials. Thus, spike analysis for determination of activation thresholds did not require filtering. For conduction block, the transmembrane potential could be driven beyond 0 mV (or another arbitrarily defined threshold) by kHz waveforms at high amplitudes. These high amplitude oscillations are “real” and not artifacts, making it difficult to distinguish action potentials which ride on top of or are merged with these oscillations. Spikes could nevertheless be identified by temporal or frequency features, because the transmembrane potential oscillations were sinusoidal and removed by filtering, while spikes resulted in distortions that remained after filtering.

In experimental studies of neural stimulation, electrophysiological recordings are contaminated by stimulus artifacts, e.g., the kHz extracellular E-field in TIS. Not only is filtering often necessary as a post-processing step, multiple filters are integral components of the experimental rig before the data are sampled [63]. In [16], the current clamp traces were band-stop filtered (1 kHz and 15 kHz) and high-pass filtered (100 Hz). While the neurons themselves were unlikely to fire at frequencies over 100 Hz in response to stimulation, it was unclear whether the filtering resulted in the loss of some spikes during kHz frequency stimulation and TIS. It appears that spikes could be identified from TIS and HFS only when the artifact amplitude was relatively small and spikes were visible even in the unfiltered traces, i.e., during the initial ramp-up of both TIS and HFS and during each ramp-up of the low frequency envelope of the TIS waveform [16]. During the steady state of HFS and near the peak of the low frequency envelope of TIS, spikes that were overwhelmed by the artifact may have been lost due to filtering, leading to a conclusion that there was no response to HFS. As such, alternate artifact removal strategies could be pursued, such as subtraction of an artifact template, or using an ideal notch filter in the frequency domain. Neuronal models are not limited by such artifacts and allow direct access to the transmembrane potential. While advantageous for data analysis, filtering of simulated transmembrane potentials should be performed with caution as it complicates the interpretation of modeling results and their comparison with experimental studies.

## 4.3 Limitations

Our simulations examined the neuronal response to uniform E-fields under the quasi-uniform approximation for transcranial stimulation. Due to this simplification, the thresholds could not be directly correlated with current amplitudes applied to tACS electrodes, and the spatial distribution of neural activation within the cortex was not determined. Nevertheless, simulations of E-field distributions in realistic head models can fill this gap, whether using a multiscale model with neuron models embedded in the cortex [32] or using the uniform field results as a lookup table to determine thresholds based on the amplitude ratio and orientations of the cortical E-fields relative to the normal direction of the cortical layer [42,64]. Therefore, the combinations of E-field orientation and their amplitude ratios in this study are representative of neurons at different cortical locations.

Despite their realistic morphologies and complexity of ion channels, the model neurons have limitations capturing the response to high frequency stimulation. The Blue Brain neuron models have truncated main axons due to the slicing process and do not include the descending axon in the white matter. Mid-axon activation of white matter fibers—thought to generate direct waves in TES at high amplitudes—could contribute to the response to TIS [25]. However, such responses have similar patterns and characteristics compared to those observed in our study and would occur at higher amplitudes as the axon terminals are the lowest-threshold sites of action potential initiation due to their sensitivity to exogenous E-fields [43,53]. Further, channel conductances and dynamics are not readily available for the axonal arbor beyond the initial segment and have to be inferred from the parameters of the initial segment. This may contribute to simulated transcranial stimulation thresholds being on average about two fold higher than experimental data, although the simulated and experimental threshold distributions overlap [32]. The sodium channel dynamics were fitted to recordings from patch clamp recording [65], which could underestimate the kinetics of axonal sodium channels due to filtering (2 kHz low pass filter) and recording below body temperature (24°C). We accelerated the fast sodium channels that contribute the most to action potential firing to capture better the response to kHz stimulation; however, further improvements are required to increase the accuracy of cortical neuron models. Nevertheless, analyses of transcranial stimulation should use cortical neuron models with realistic myelinated axonal morphology at body temperature, and classic Hodgkin-Huxley channels [66,67] are ill suited to studies of transcranial stimulation in mammals.

We did not include intrinsic activity or network connectivity in this study. Although patch clamp recordings in the anesthetized mice exhibited negligible spontaneous firing (< 0.5 Hz) [16], intrinsic subthreshold fluctuation of the transmembrane potential could have interacted with the exogenous E-field. However, threshold changes for stimulation during the afterpolarization of an action potential are on the order of ±50% [68], and therefore transmembrane potential fluctuations alone are unlikely to account for the discrepancy of threshold differences between LFS and kHz tACS when comparing the experiments and our simulations.

As our goal was to explore the hypothesis of single-neuron demodulation of TIS, we did not include synaptic connections between neurons, and network modulation could be a potential mechanism for TIS. While network activation can be advantageous, it could also be considered a limiting factor on the focality of transcranial stimulation methods, including TIS. For example, Grossman et al. [16] used c-Fos immunochemistry to demonstrate activation of the hippocampus without activating superficial cortical areas, showing strong c-Fos immunoreactivity in the entire region of granule cells within the dentate gyrus. This expression was much stronger than observed anywhere for LFS or elsewhere in TIS, suggesting that TIS achieved strong synchronous activation of the local circuitry in the dentate gyrus that could potentially be a result of a focal seizure. Furthermore, the area of activation inferred from the immunohistochemistry does not correlate with the distribution of the simulated E-field modulation envelope amplitude [16], complicating the interpretation of the mechanism and focality of TIS. While these limitations are important directions for future refinement of the models, they would affect similarly TIS and the other forms of kHz stimulation. Therefore, our conclusions regarding the relative effects and thresholds across these modalities appear unlikely to be altered.

## 5. Conclusions

We analyzed the E-field characteristics of TIS and quantified the responses of model cortical neurons to suprathreshold TIS and related kHz tACS modalities. The analysis revealed unique spatiotemporal characteristics of the TIS E-field, including phase-shift and frequency jitter of the TIS waveforms due to the difference in E-field amplitudes for aligned E-fields, as well as rotation of the total E-field for non-aligned E-fields. These findings highlight the importance of representing the TIS E-field in 3D and using morphologically-realistic neuron models. TIS had activation thresholds similar to other kHz tACS modalities and much higher than those of low frequency tACS. As expected, the bursts of activation evoked by near-threshold TIS were synchronized with the maxima of the low frequency modulation envelope of the total E-field, providing one possible mechanism for the previously reported experimental responses. Further, at amplitudes exceeding the activation threshold by an order of magnitude, TIS generated neuronal block, and in some cases the modulation of the total E-field generated periodic onset responses, providing an alternative mechanism for the experimental observations. Conduction block may also explain the lack of responses in off-target regions, where block can suppress endogenous brain activity. While the neuron models may overestimate absolute activation thresholds, the E-field thresholds for activation and block with TIS are, respectively, two and three orders of magnitude higher than the field strength than can be generated with tolerable levels of scalp stimulation. Therefore, administering suprathreshold TIS with transcranial electrical stimulation of the human brain appears infeasible without anesthesia and any TIS application should consider off-target effects of the carrier.

## Acknowledgments

This work was supported by grants No. R01NS088674 and No. R01NS117405 from the National Institutes of Health of the United States of America; the content is solely the responsibility of the authors and does not necessarily represent the official views of the funding agency or U.S.A. government. Computational support was provided by the Duke Compute Cluster.

The authors thank Dr. Nir Grossman for providing critical feedback on preliminary results at the 2^nd^ Carolina Neuromodulation Conference, Dr. Flavio Fröhlich for discussion on activation mechanism of TIS, Dr. Nicole A Pelot for discussion on onset response of kHz stimulation, Dr. Craig S. Henriquez and Dr. Derek M. Eidum for discussion on rotational E-fields, Dr. Sumientra Rampersad for discussion on E-field simulation in mouse model, and Dr. Huijing Xu, Dr. Gene Yu, and Dr. Alex Goddard for discussion on stimulation of hippocampal neurons.

B. W. developed the theoretical framework, designed the computational study, and performed the neural simulations and data analysis. A. S. A. developed the neuronal models and adjusted ion channel properties. W. M. G. and A.V.P. conceived, supervised, and secured funding and computational resources for the study. B. W. performed data visualization and wrote the manuscript, and all authors revised, commented on, and approved the final version of the manuscript.

Preliminary results of this study were presented at the 1st and 2nd Carolina Neuromodulation Conference (May 2018 and June 2019, Chapel Hill, NC, USA), the 43rd Neural Interface Conference (June 2018, Minneapolis, MN, USA), Neural Interfaces 2021: The NANS-NIC Joint Meeting (June 2021, online), and the 4th International Brain Stimulation Meeting [69] (December, 2021, Charleston, SC, USA).

The code and data that support the findings of this study are available online at the Research Data Repository (RDR) of Duke University Libraries: https://doi.org/10.7924/r4n87d05r.

The authors declare no competing interests.

## Supplement

**Figure S1.**
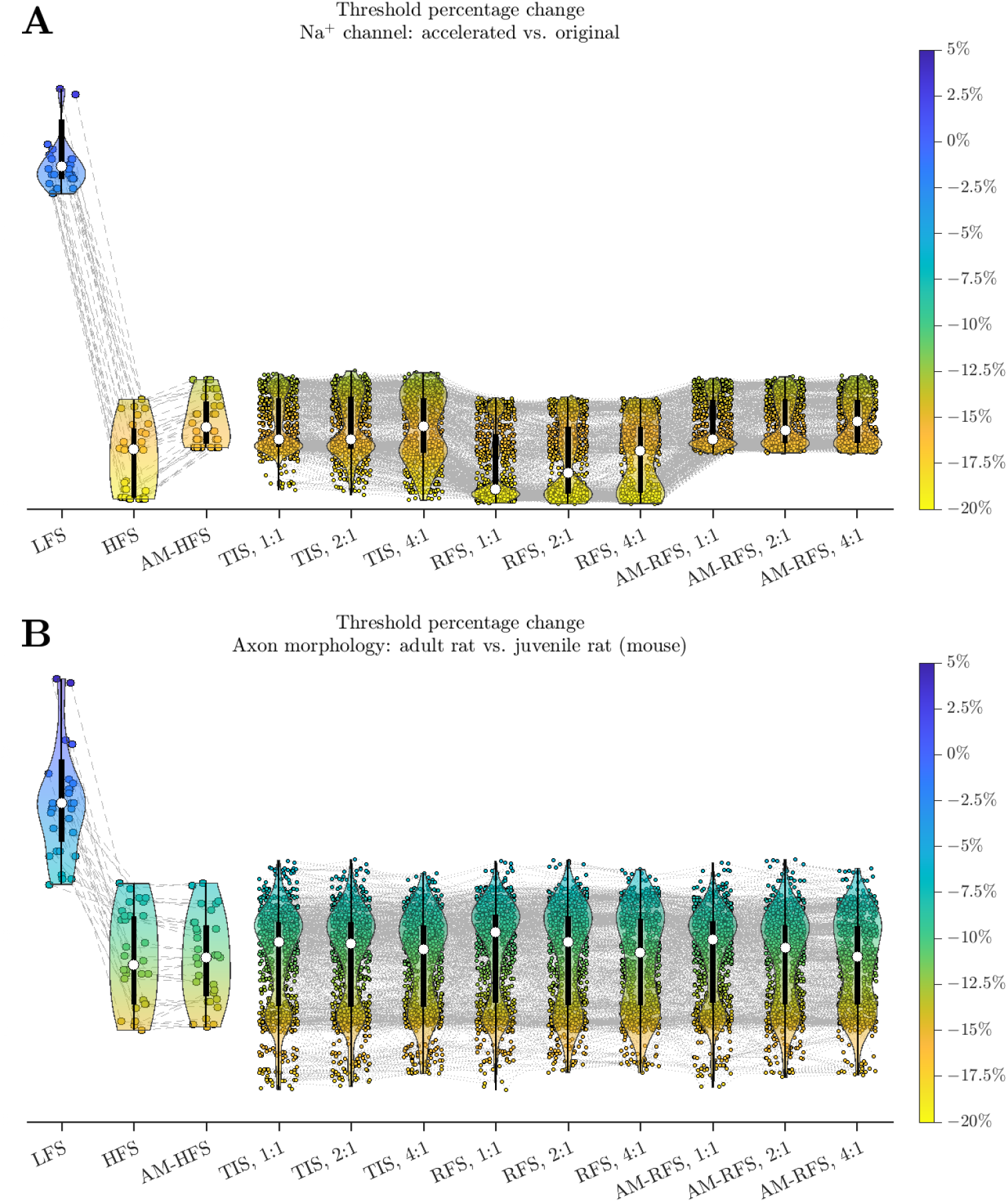
Changes of activation thresholds for modification applied to the neuron models. **A**. Percentage changes of thresholds of the tACS modalities across E-field orientations comparing accelerated sodium channel dynamics versus original. **B**. Percentage changes of thresholds of the tACS modalities across E-field orientations for neuron models scaled to size of adult rat from original size of juvenile rat. The latter size is used to simulate adult mice in this study. The limited change of thresholds demonstrate that the size difference between rat and mouse neurons is not critical for comparing the neuronal response to the types of stimulation in this study. Figure format is overall similar to Figure 7. Outliers of LFS were excluded. The gray dashed lines connect the same E-field orientations between the static modalities and orientation combinations between the non-static modalities.

**Figure S2.**
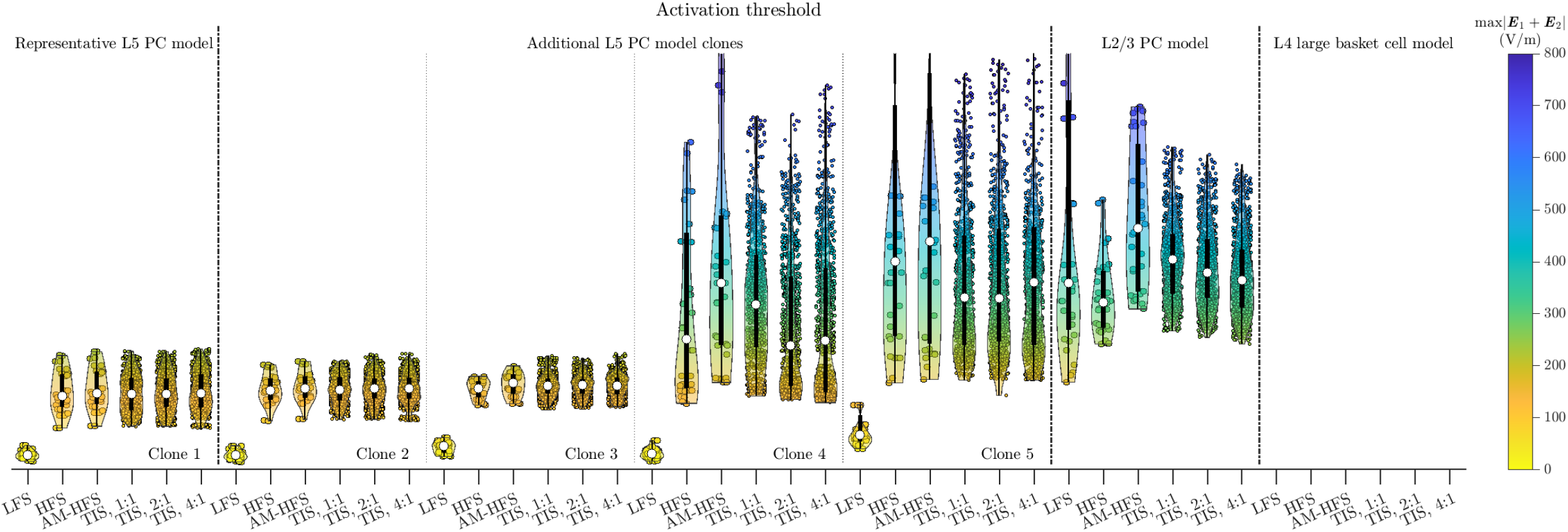
Activation thresholds of LFS, HFS, AM-HFS, and TIS for five model clones of the layer 5 pyramidal cell (L5 PC), one layer 2/3 PC, and one layer 4 interneuron. Figure format is overall similar to Figure 7, with the results for each clone separated by vertical dashed lines. The representative L5 model (clone No. 1) used is shown on the left. Two L5 clones (No. 2 and 3) have threshold distributions similar to the representative model, whereas the other two clones have wider threshold distributions for kHz tACS due to their axon morphologies. For the latter two L5 clones (No. 4 and 5), the ratio of the lowest kHz thresholds versus LFS thresholds is comparable to those of the first three clones. Moreover, the additional clones do not exhibit lower absolute thresholds than the representative model. Therefore, the conclusions in the main text based on the representative L5 model hold generally. The L2/3 PC also has higher threshold for all stimulation modalities than the representative L5 model, and the LFS threshold had a wide range due to the model neuron lacking axon terminals aligned with some orientations of the E-field. Due to its dense dendritic morphology and different ion channel conductivities, the L4 interneuron had thresholds ≫ 1 kV/m for a few E-field orientations whereas for the majority of E-field orientations it could not be activated periodically at 10 Hz by E-field of strength in the search range up to 4000 V/m. Therefore the results of L4 interneuron are not visible in the figure.

**Figure S3.**
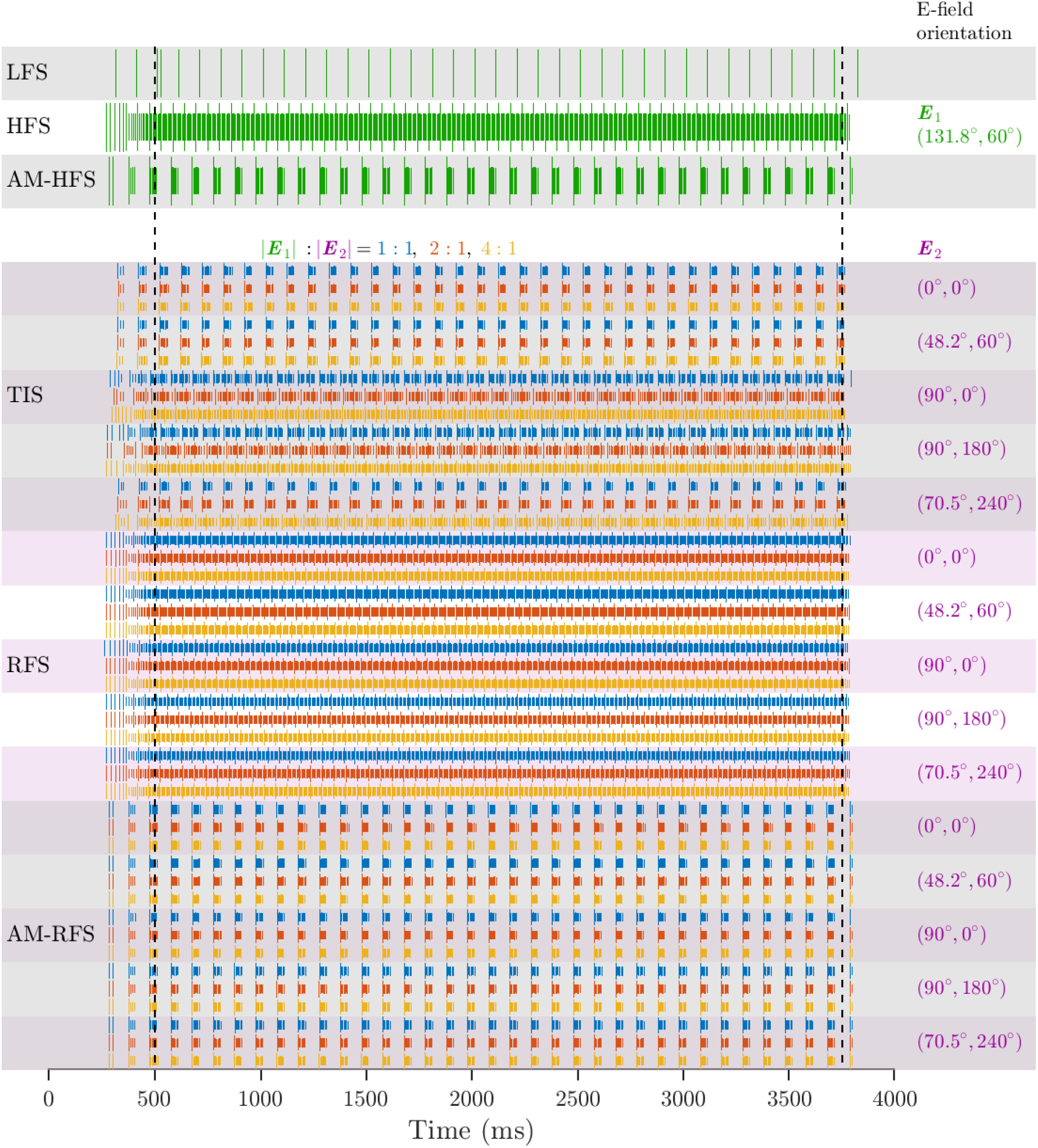
Representative suprathreshold responses of L5 PC. Raster plot of somatic firing for stimulation amplitude at 1.5 times activation threshold as shown in Figure 6. The first spike within each burst is shown with a longer line.

**Figure S4.**
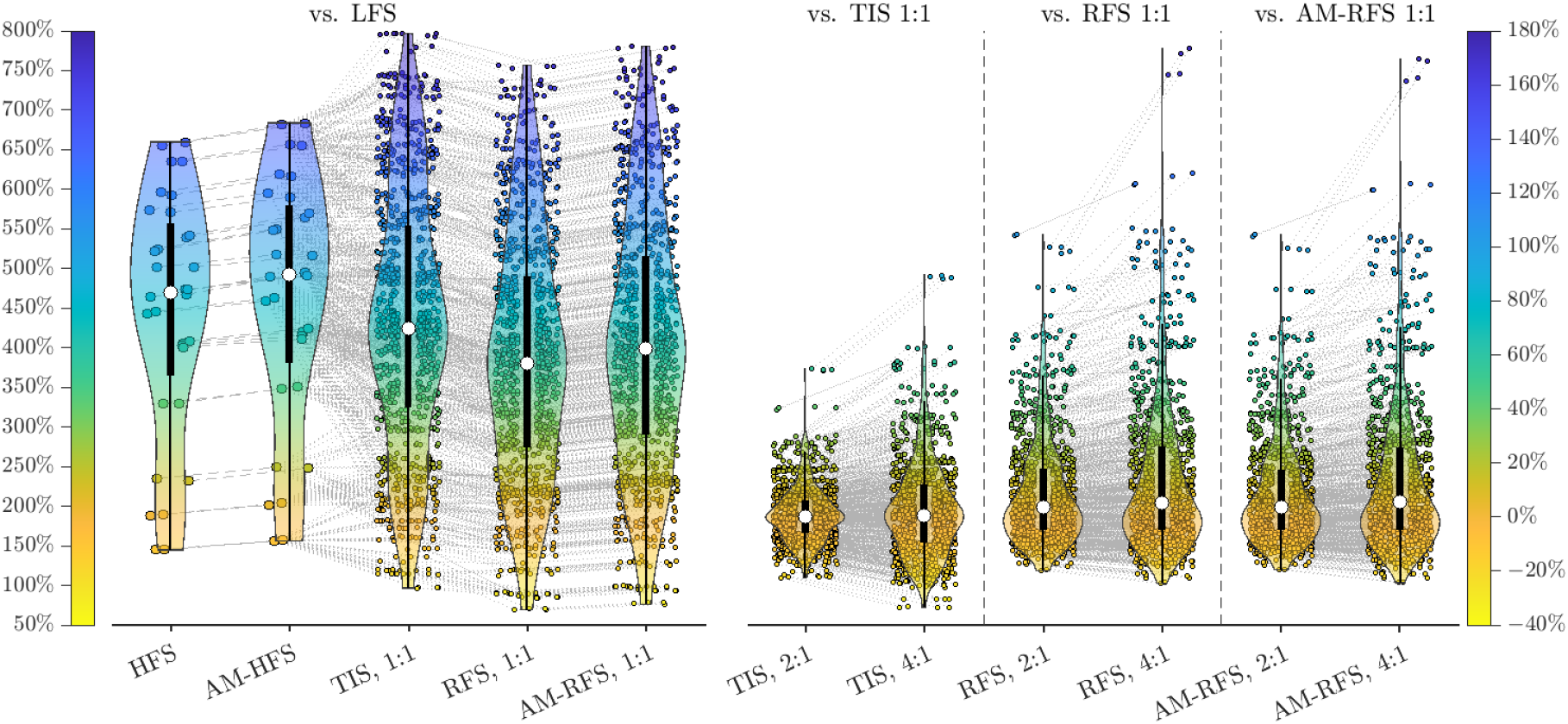
Changes of activation thresholds. Left: percentage changes of thresholds versus LFS for HFS and AM-HFS, as well as TIS, RFS, and AM-RFS of equal E-field amplitudes. For HFS and AM-HFS, the changes were calculated against LFS with the same E-field orientation. For TIS, RFS, and AM-RFS, the comparison was against LFS with the E-field orientation aligning closest with the E-field orientation at maximum amplitude. The gray dashed lines connect the same E-field orientations between HFS and AM-HFS and orientation combinations between the non-static modalities, and links the E-field orientation of the static stimulation modalities with the closest non-static E-field at maximum amplitude. Right: percentage changes of thresholds of non-static tACS modalities with unequal E-field versus equal E-field. Figure format is similar to Figure 7.

**Figure S5.**
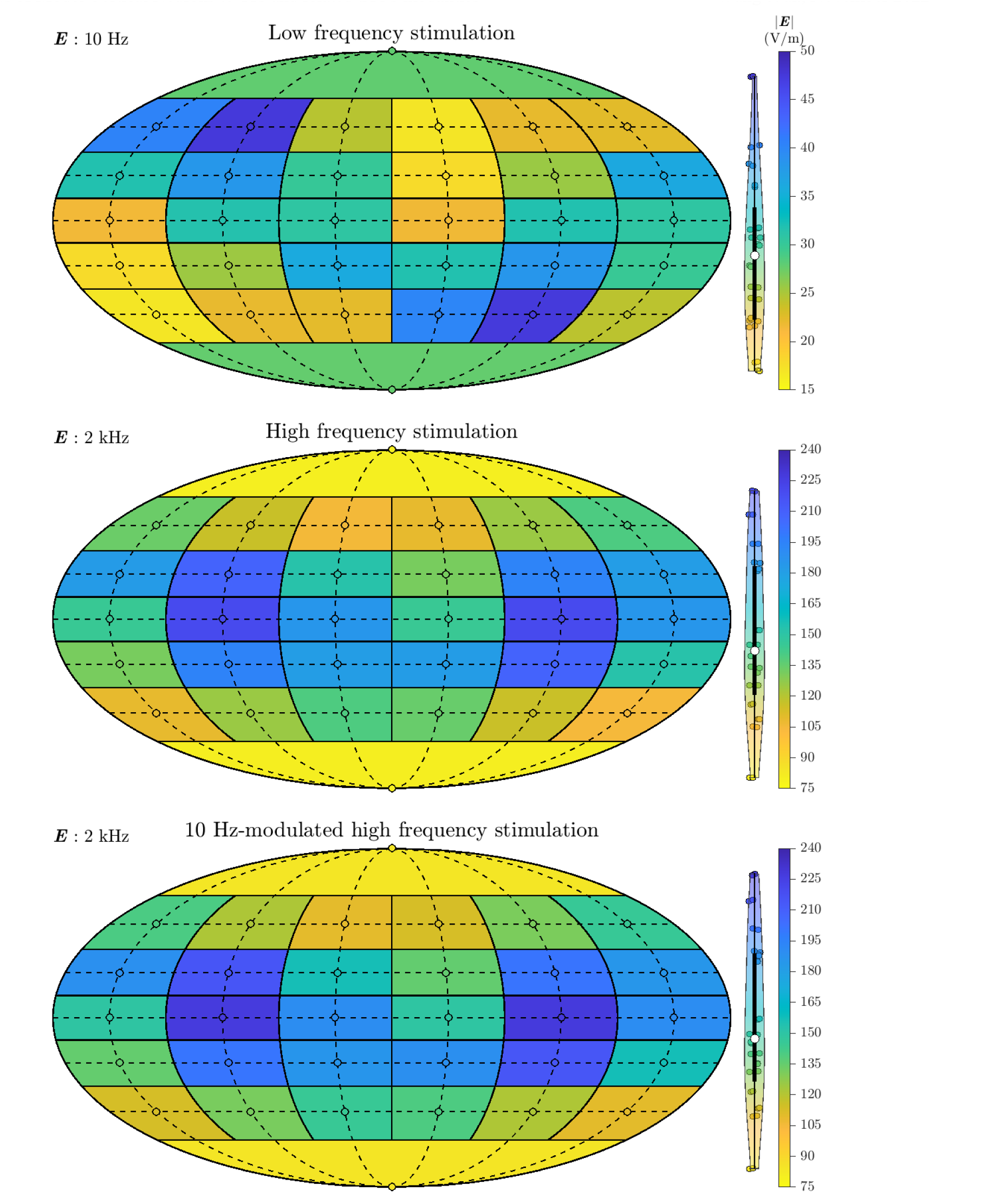
Distributions of activation threshold for all E-field orientations of LFS, HFS, AM-HFS. The orientations of the E-field is converted from its spherical arguments into 2D coordinates with Mollweide projection. Each patch shows the threshold of one orientation (circle) with dashed line showing the grid of the E-field sampling in the spherical coordinate.

**Figure S6.**
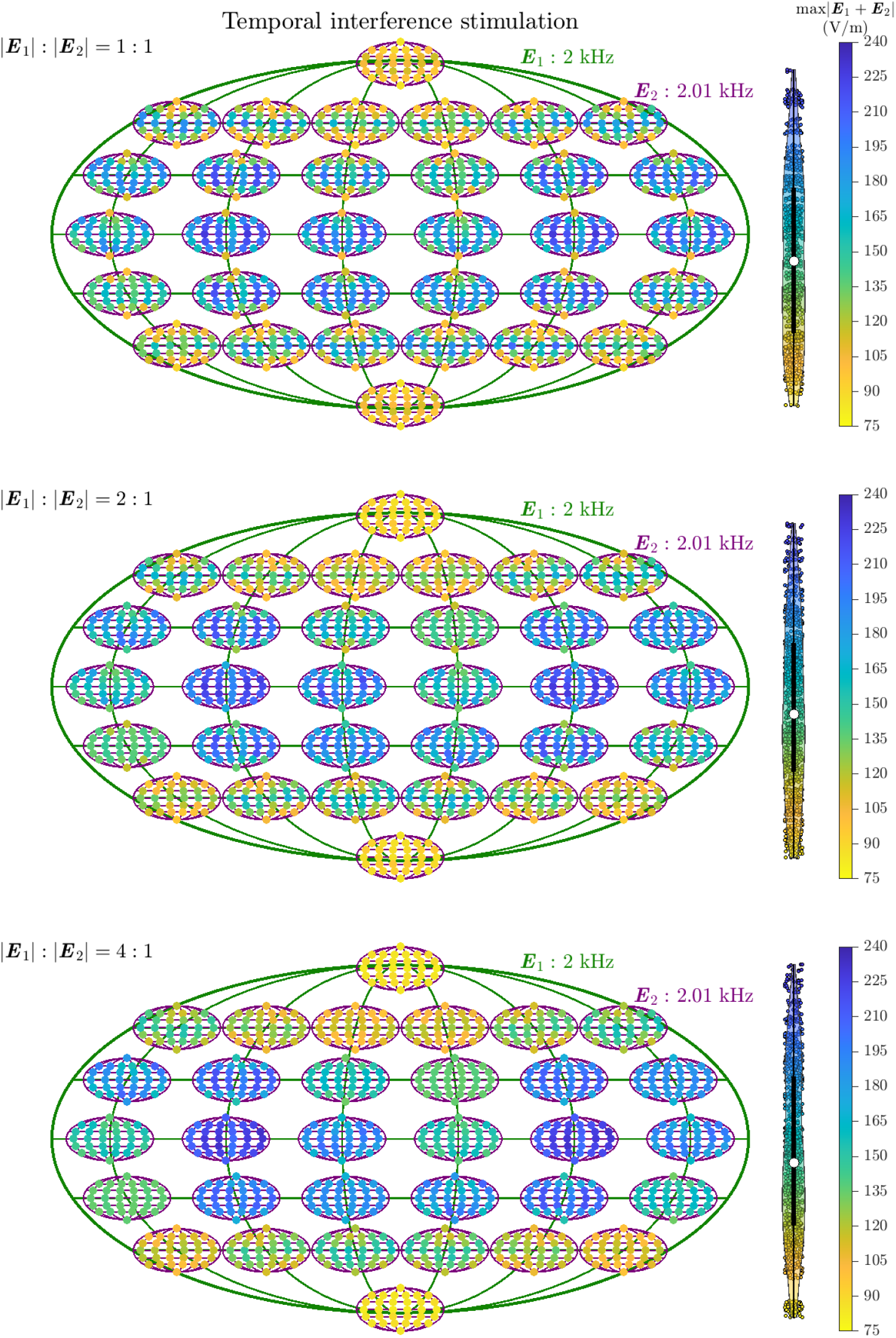
Distributions of activation threshold for all E-field orientation combinations of TIS. Both E-fields ***E***_1,2_ are mapped using Mollweide projection, with the smaller maps showing the thresholds for all 32 orientations of the second E-field ***E***_2_ for a fixed orientation of the first E-field ***E***_1_, and the smaller maps are arranged according to the orientations of ***E***_1_.

**Figure S7.**
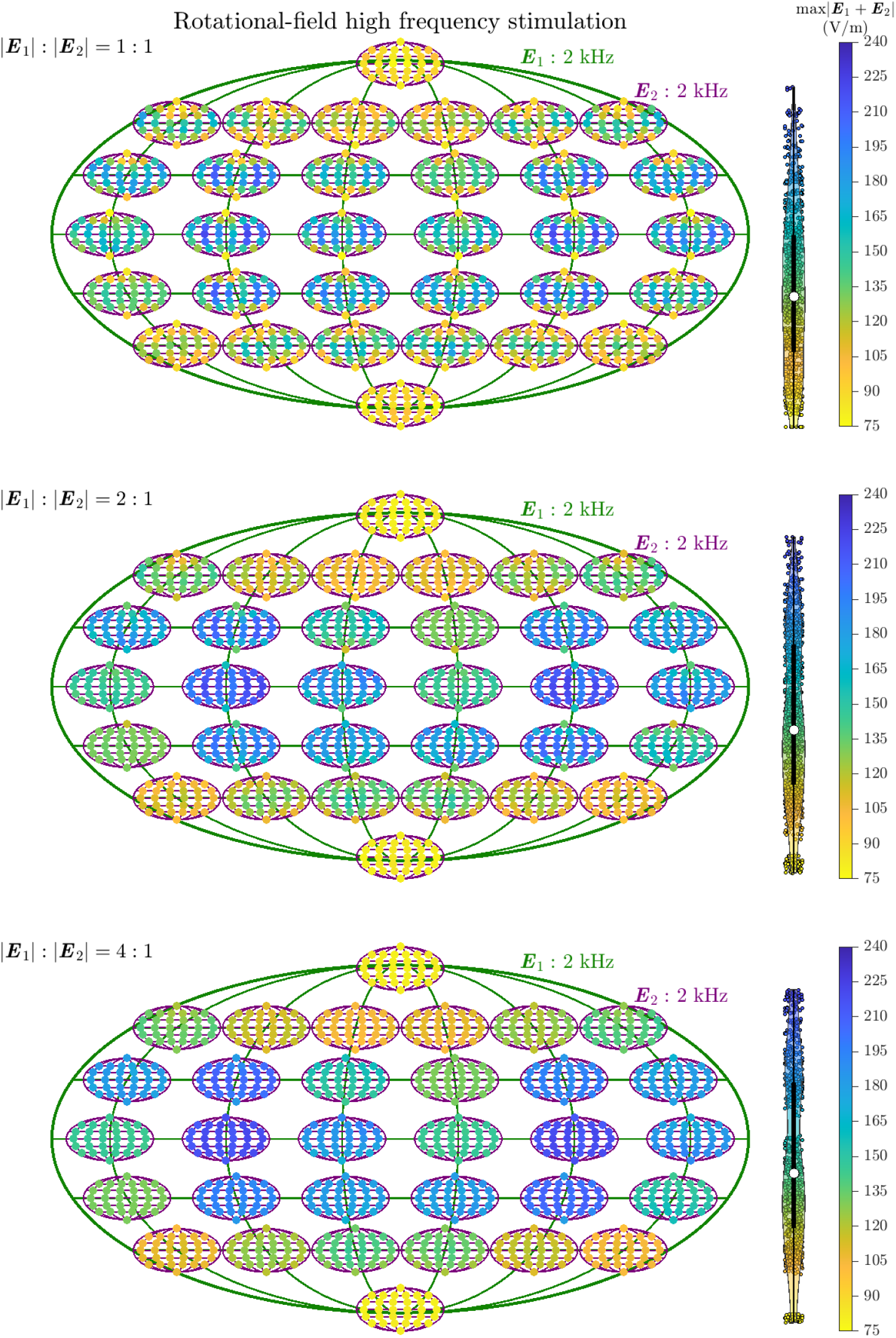
Distributions of activation threshold for all E-field orientation combinations of RFS. Same format as Figure S6.

**Figure S8.**
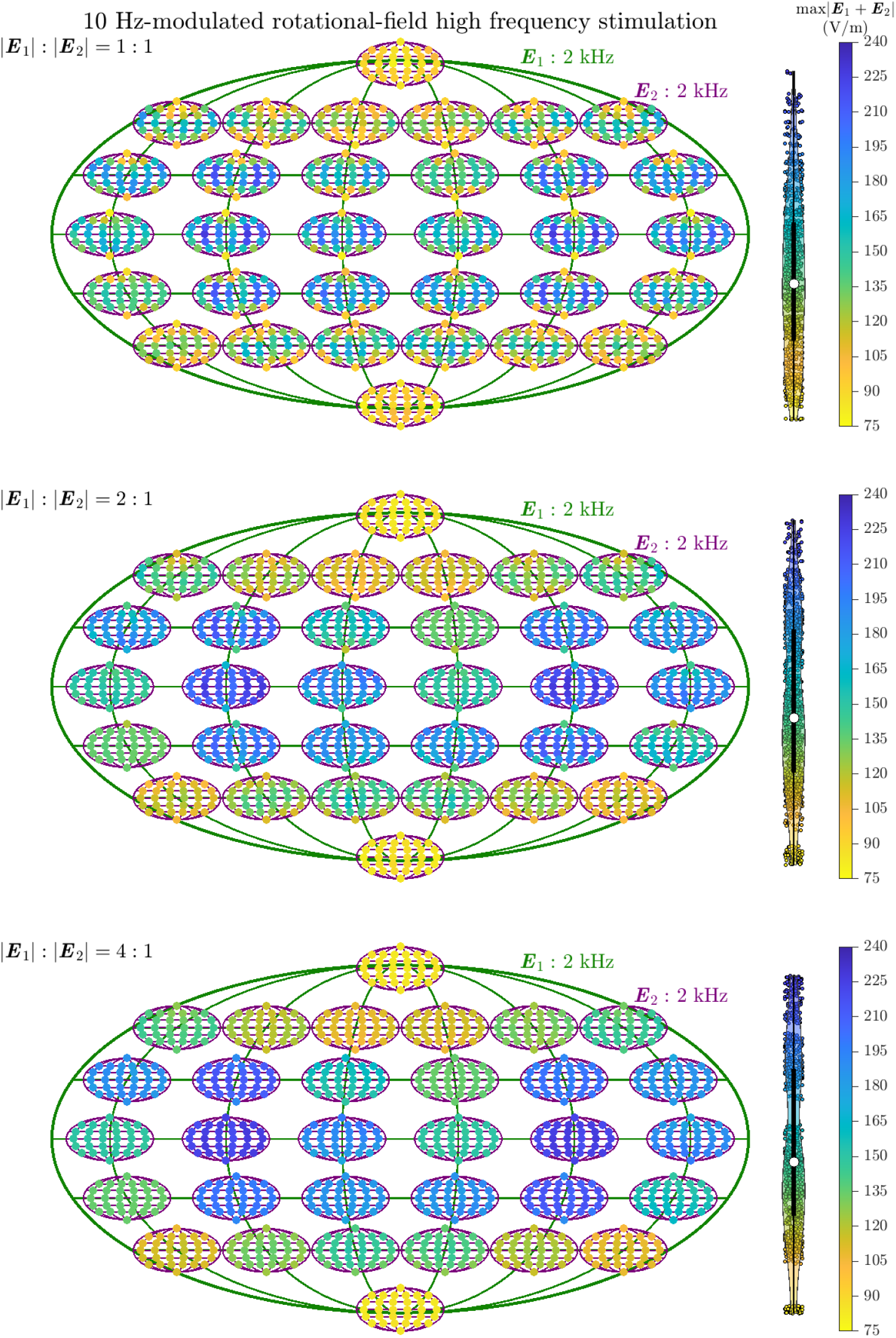
Distributions of activation threshold for all E-field orientation combinations of AM-RFS. Same format as Figure S6.

**Figure S9.**
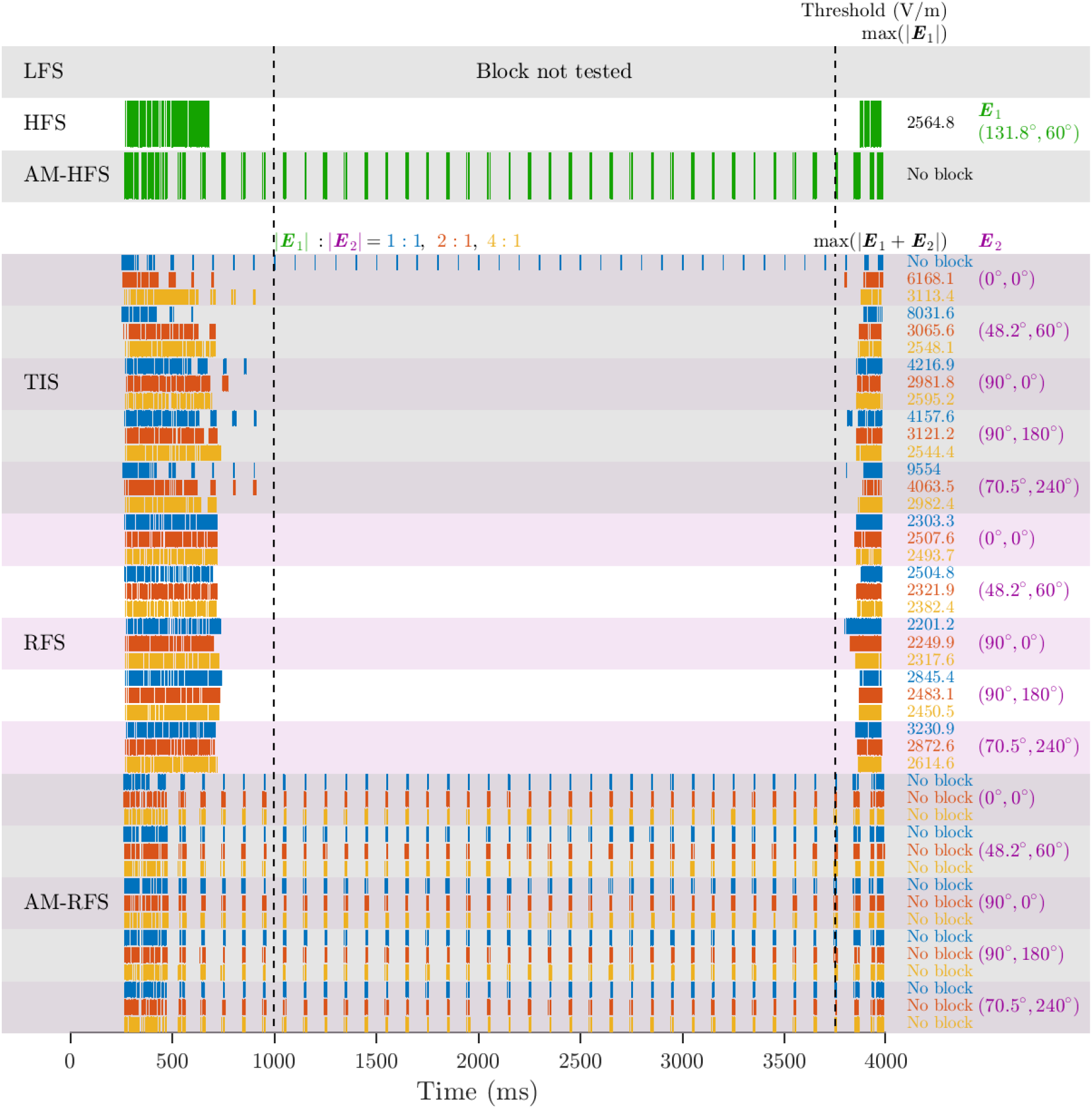
Representative responses of L5 PC at conduction block thresholds or maximum amplitude tested. Same as Figure 8, but for spikes of the raw transmembrane potential.

**Figure S10.**
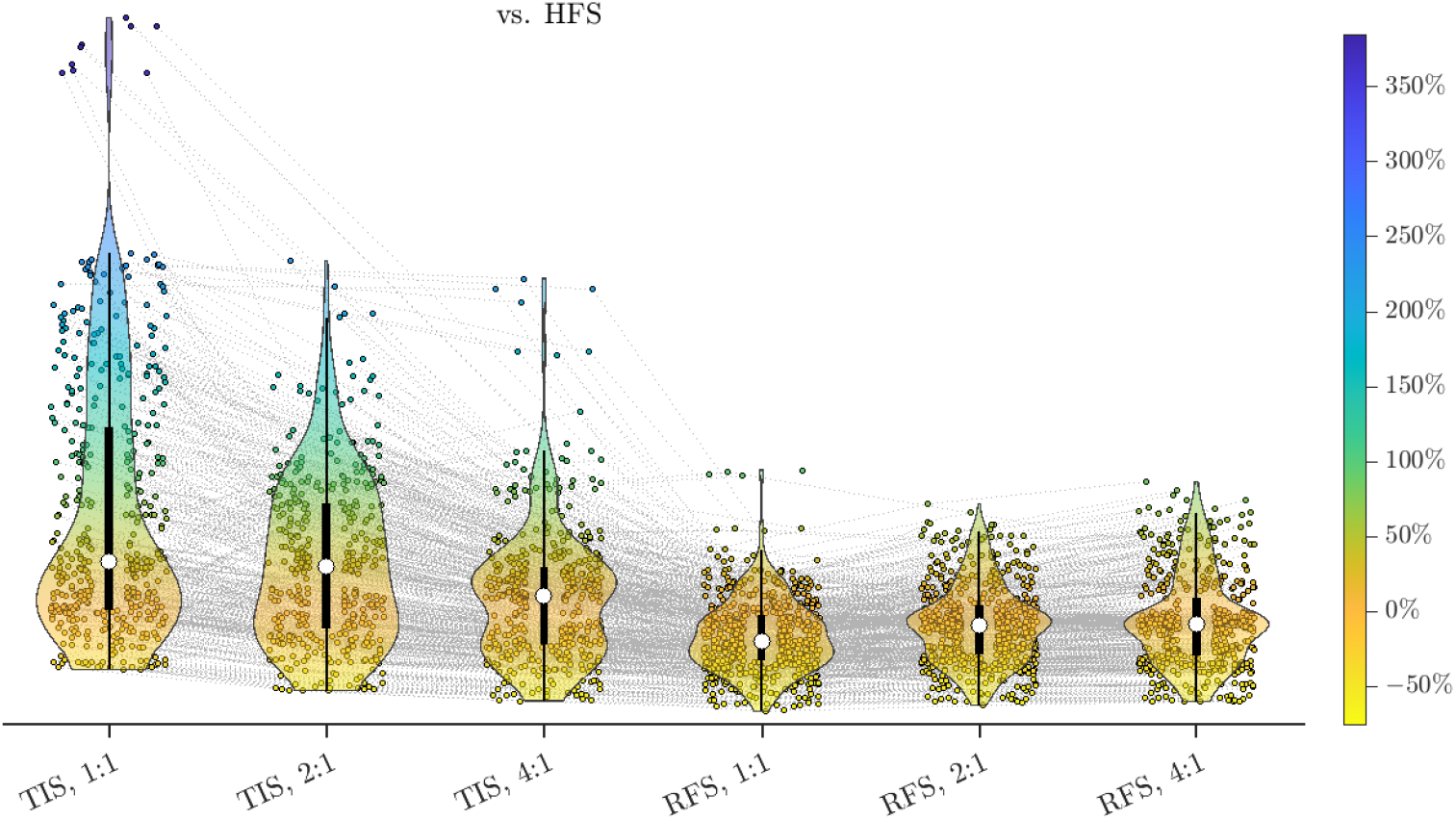
Changes of conduction block thresholds. Percentage changes of TIS and RFS thresholds versus HFS, calculate against HFS with E-field orientation aligning closest with the TIS or RFS E-field orientation at maximum amplitude. The gray dashed lines connect the same E-field orientation combinations between the non-static modalities. Similar format as Figure S4.

**Figure S11.**
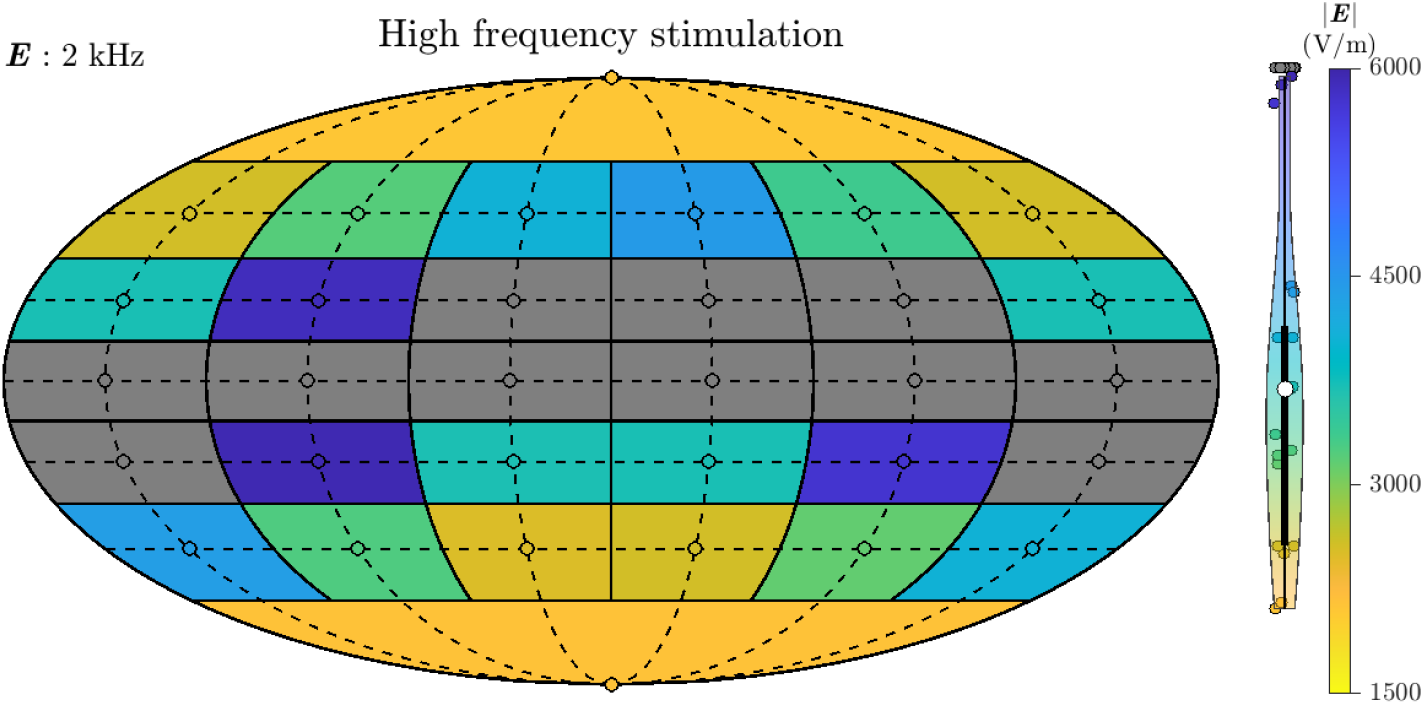
Distributions of conduction block threshold for all E-field orientations of HFS. Similar format as Figure S5. Grey data points indicate that conduction block could not be achieved within 6000 V/m.

**Figure S12.**
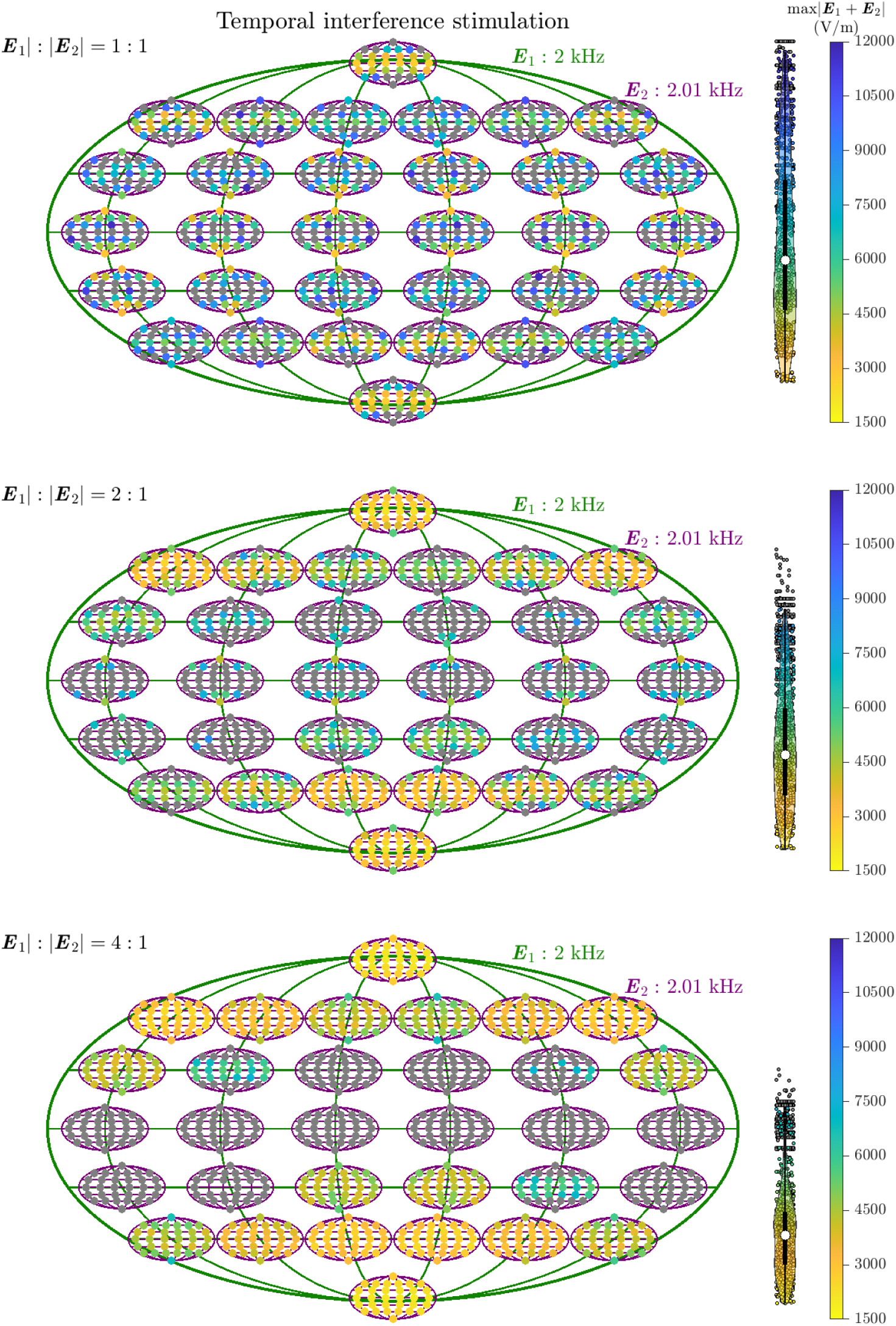
Distributions of conduction block threshold for all E-field orientation combinations of TIS. Similar format as Figure S6. Gray data points indicate that block could not be achieved within 6000 V/m.

**Figure S13.**
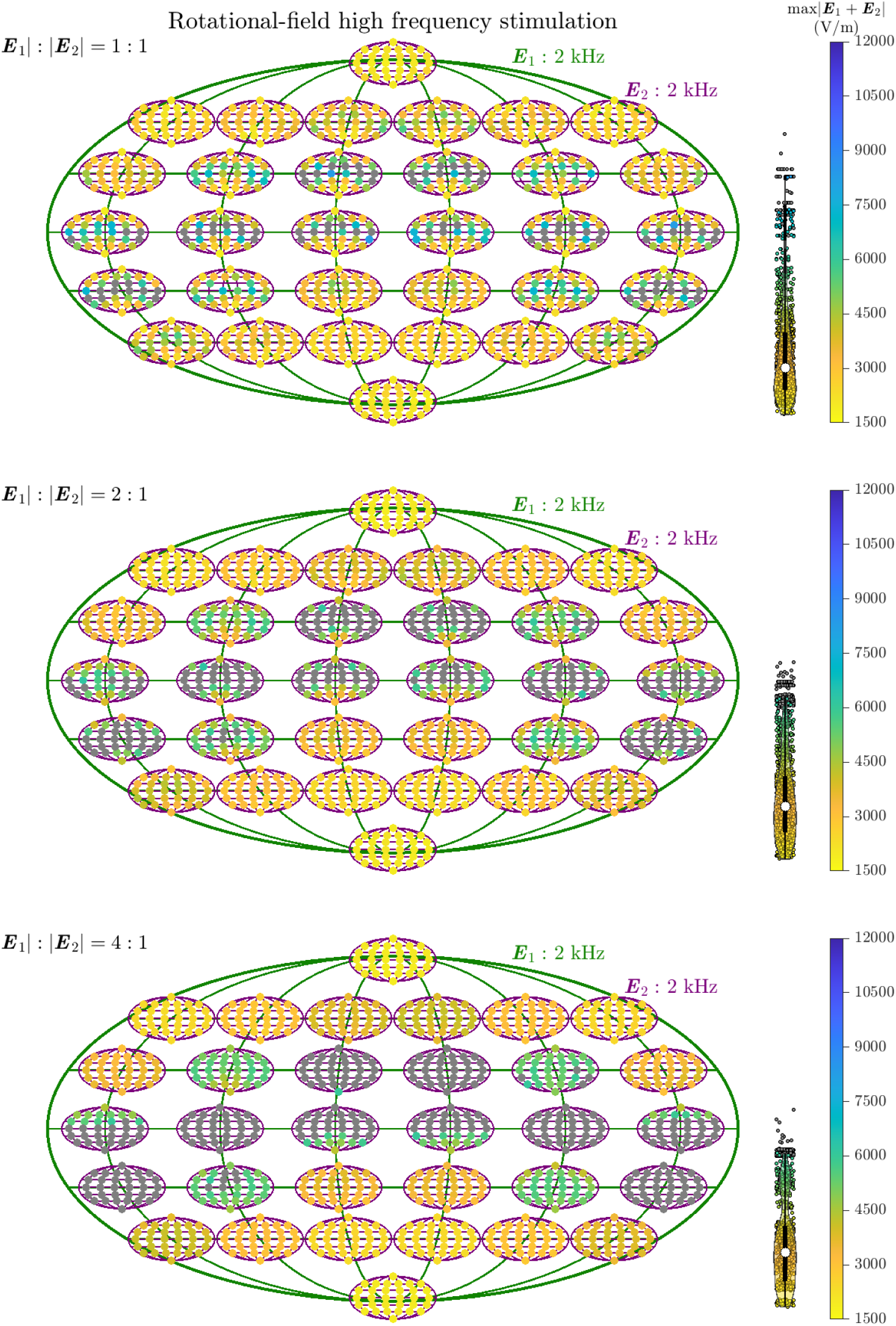
Distributions of conduction block threshold for all E-field orientation combinations of RFS. Same format as Figure S12.

**Figure S14.**
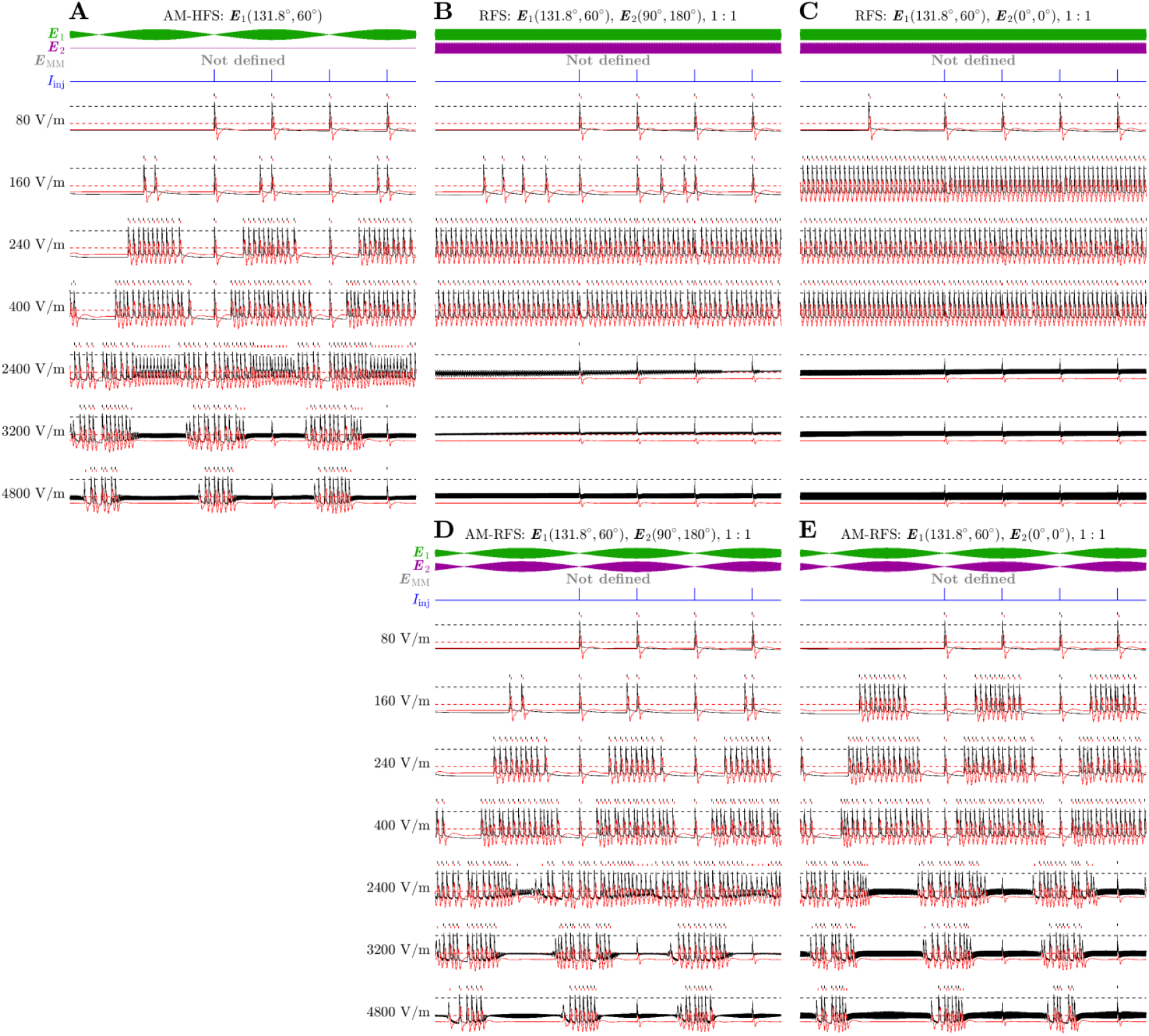
Neuron firing evoked by AM-HFS, RFS and AM-RFS for a range of representative E-field amplitudes. Similar format as Figure 9. E-field amplitudes are given as either max(|***E***_1_|) for AM-HFS or max(|***E***_1_+***E***_2_|) for RFS and AM-RFS. **A**. AM-HFS evokes periodic firing, which shifts from the maximum of the envelope to the minimum for high stimulation amplitude, indicating that the firing becomes an onset/offset response. Test pulses injected at the maximum of the envelope are blocked. **B–C**. RFS results in sustained firing that is blocked at higher stimulation amplitudes. **D–E**. AM-RFS results in periodic activation that shifts from the maximum of the envelope to the minimum like AM-HFS. The test pulses at the maximum of the envelope are blocked.

